# Self-assembly of multi-component mitochondrial nucleoids via phase separation

**DOI:** 10.1101/822858

**Authors:** Marina Feric, Tyler G. Demarest, Jane Tian, Deborah L. Croteau, Vilhelm A. Bohr, Tom Misteli

## Abstract

Mitochondria contain an autonomous and spatially segregated genome. The organizational unit of their genome is the nucleoid, which consists of mitochondrial DNA (mtDNA) and associated architectural proteins. Here, we show that phase separation is the primary physical mechanism for assembly and size-control of the mitochondrial nucleoid. The major mtDNA-binding protein TFAM spontaneously phase separates *in vitro* via weak, multivalent interactions into viscoelastic droplets with slow internal dynamics. In combination, TFAM and mtDNA form multiphase, gel-like structures *in vitro*, which recapitulate the *in vivo* dynamic behavior of mt-nucleoids. Enlarged, phase-separated, yet transcriptionally active, nucleoids are present in mitochondria from patients with the premature aging disorder Hutchinson-Gilford Progeria Syndrome (HGPS) and are associated with mitochondrial dysfunction. These results point to phase separation as an evolutionarily conserved mechanism of genome organization.

**Highlights:**

- Mitochondrial genomes are organized by phase separation.
- The main packaging protein TFAM and mtDNA combine to form viscoelastic, multiphase droplets *in vitro*.
- Mitochondrial nucleoids exhibit phase behavior *in vivo*, including dynamic rearrangements and heterogenous organization.
- Coalescence and enlargement of mt-nucleoids occur upon loss of mitochondrial homeostasis as well as in prematurely aged cells and are associated with mitochondrial dysfunction.

## Introduction

Mitochondria are the major sites of cellular energy production through oxidative phosphorylation and generation of ATP. Within each mammalian cell, mitochondria contain several hundred copies of a 16.6 kb double-stranded circular genome (Chen and Butow, 2005; Gustafsson et al., 2016). Although small in size, the mitochondrial genome is gene dense, encoding essential polypeptides involved in mitochondrial respiration and oxidative phosphorylation (Burger et al., 2003). Unlike the nuclear genome, mitochondrial DNA (mtDNA) is not organized by histones, but is packaged by a distinct set of proteins into nucleoprotein complexes to form mitochondrial nucleoids (mt-nucleoids) (Chen and Butow, 2005; Kukat and Larsson, 2013). These structures are typically uniformly ∼100 nm in size, each containing 1-2 molecules of mtDNA, and are spatially separated throughout the mitochondrial network (Brown et al., 2011; Kukat et al., 2011). mt-nucleoids lack any delimiting membranes, yet act as discrete functional units involved in replication and transcription of the mitochondrial genome (Kukat and Larsson, 2013).

The major mt-nucleoid packaging protein in human cells is the mitochondrial transcription factor A (TFAM) (Chen and Butow, 2005), which binds and compacts mtDNA *in vitro* into nucleoid-like structures under dilute conditions (Brewer et al., 2003; Farge et al., 2014; Kaufman et al., 2007; Kukat et al., 2015). TFAM contains two high mobility group (HMG) domains that each intercalate into the DNA double helix, bending the DNA strand to form a tight U-turn structure at promoter sequences (Ngo et al., 2011; Rubio-Cosials et al., 2011). Moreover, TFAM can also form loops and bind cross strands of DNA without sequence specificity (Kukat et al., 2015), and these conformations are further stabilized by cooperative TFAM-TFAM interactions (Farge et al., 2012). In human cells, TFAM is present at high enough concentrations to coat the entirety of the circular mtDNA, driving the compaction of mtDNA from ∼5 um in contour length to the ∼100 nm mt-nucleoid (Gustafsson et al., 2016). Beyond the direct binding of TFAM to mtDNA, it remains largely unknown how the higher-order morphological features of the mitochondrial genome emerge, how they affect function, and how anomalies in the structure of mt-nucleoids may contribute to disease (Friedman and Nunnari, 2014).

Maintenance of mt-nucleoid structure is linked to mitochondrial organization and function. Disruption of mitochondrial fusion and fission processes affect nucleoid size, as seen in knockdown of mitochondrial fission GTPase, Drp1, which leads to clustering of nucleoids into large assemblages in hyperfused mitochondria (Ban-Ishihara et al., 2013; Ishihara et al., 2015). Similarly, downregulation of the inner mitochondrial membrane protein Mic60/Mitofilin leads to disassembly of the mitochondrial contact site and cristae organizing system (MICOS), ultimately resulting in enlarged, spherical mt-nucleoids (Li et al., 2016). The enlargement and remodeling of mt-nucleoids are also associated with a cellular response to stress. For example, prolonged exposure to DNA intercalating agents leads to altered nucleoid size distributions (Alán et al., 2016; Ashley and Poulton, 2009), and viral infection results in aberrant sizes of mt-nucleoids (West et al., 2015). Given the oxidative environment within mitochondria (Balaban et al., 2005; Sun et al., 2016) and the absence of protective histone proteins, the nucleoid associated proteins have been hypothesized to contribute to mitochondrial genome integrity (Cadenas and Davies, 2000; Yakes and Van Houten, 1997). Mutations in mtDNA have direct physiological relevance as elevated mutation levels of mtDNA are associated with premature aging phenotypes (Kujoth et al., 2005; Trifunovic et al., 2004). mtDNA mutations tend to accumulate over the course of normal aging (Bratic and Larsson, 2013; Sun et al., 2016) and even single point mutations in mtDNA can elicit a myriad of other disease phenotypes (Taylor and Turnbull, 2005).

An emerging organizational principle of non-membrane bound cellular structures is phase separation (Hyman et al., 2014). Numerous ribonucleoprotein and nucleoprotein complexes spontaneously self-assemble into non-membrane bound cellular bodies, or biomolecular condensates, via liquid-liquid phase separation (Banani et al., 2017). The canonical examples of RNA-protein bodies include the nucleolus in the nucleus (Brangwynne et al., 2011; Feric et al., 2016) as well as P-granules (Brangwynne et al., 2009) and stress granules in the cytoplasm (Guillén-Boixet et al., 2020; Molliex et al., 2015; Sanders et al., 2020; Yang et al., 2020). In addition, DNA-protein complexes can phase separate in the nucleus, such as the heterochromatin protein HP1α in the context of heterochromatin (Larson et al., 2017; Strom et al., 2017), histones to form chromatin domains (Gibson et al., 2019; Sanulli et al., 2019), or super-enhancers which form active transcriptional hubs (Sabari et al., 2018).

Here, we have explored the higher-order organizational principles of the mitochondrial genome in health and disease. We demonstrate, based on *in vitro* and *in vivo* observations, that mitochondrial nucleoids self-assemble via phase separation. We find that the major mt-nucleoid protein TFAM exerts its architectural role by promoting phase separation via weak, multivalent self-interactions to generate the multiphasic mt-nucleoid structure. We also demonstrate that aberrant mt-nucleoid size is associated with mitochondrial dysfunction in the context of premature aging. Our observations suggest phase separation is an evolutionarily conserved mechanism in genome organization.

## Results

### Enlarged mitochondrial nucleoids *in vivo*

During the course of in depth analysis of their cellular morphology, we noticed the presence of aberrant mitochondria and enlarged mt-nucleoids in skin fibroblasts from patients with the premature aging disorder Hutchinson-Gilford Progeria Syndrome (HGPS) (Figure 1). HGPS is a rare, invariably fatal premature aging disorder characterized by multi-tissue symptoms, including of bone, muscle, skin and cardiovascular failure. The disease is caused by a point mutation in *LMNA* resulting in the production of progerin, a dominant negative form of the major architectural protein lamin A (Gordon et al., 2014). In line with mitochondrial abnormalities associated with HGPS (Rivera-Torres et al., 2013; Xiong et al., 2016), ∼70% of advanced HGPS patient cells had a sub-population of mitochondria that were swollen, spherical in shape and isolated from the surrounding mitochondrial network compared to the typical tubular, elongated mitochondrial networks in control cells (Figures 1A, 1B, and S1E-S1K). The extent and number of enlarged mitochondria correlated with disease progression (Figures 1C and S1K) and several chaperones and proteases of the mitochondrial unfolded protein response (UPR^mt^) including HSPD1 (mtHSP60), mtHSP10, mtHSP70, ClpP, and LONP1, were enriched in enlarged mitochondria, indicating that the altered mitochondrial morphology is associated with mitochondrial stress (Figures 1J-1L and S1Q-S1V). Exogenous expression of progerin, the disease-causing isoform of lamin A in HGPS, was sufficient to induce in wild-type cells an increase in the number of aberrant mitochondria that scaled with progerin expression (Figures S1W-S1AE).

**Figure 1:**
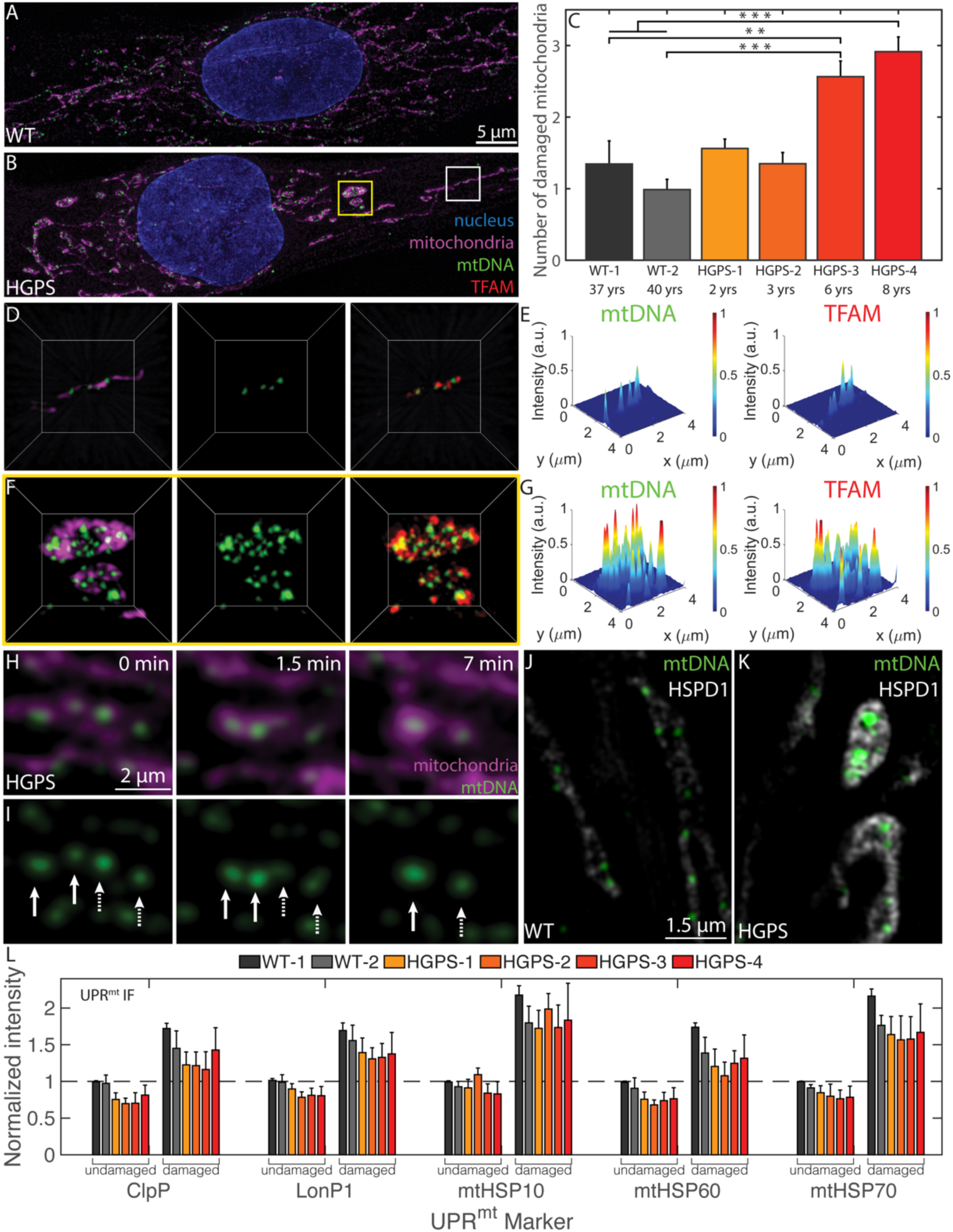
Enlarged mitochondrial nucleoids are prominent in a premature aging disease and can also arise from liquid-like fusion events under stress. (A-B), Maximum intensity projections of SIM images of a fixed normal (A) and an HGPS (B) human skin fibroblast, where the mitochondria are labelled in magenta with MitoTracker Red, mtDNA nucleoids in green with anti-DNA, and the nucleus in blue with DAPI. Scale bar = 5 µm. **c**, Bar graph quantifying the number of damaged mitochondria per cell based on high-throughput imaging of two wild-type and four HGPS primary skin fibroblast cell lines. Error bars represent averages ±SEM for n=3 experimental replicates (each experimental replicate had 15 technical replicates each containing 5 fields of view, approximately 2,000-5,000 total cells for each cell line analyzed), where p-value for the ANOVA test statistic was p<0.001. For individual pairs, **p<0.01, ***p<0.001. (D-F) Three-dimensional views of normal mitochondria (D) annotated by white box in B and swollen mitochondria (F) annotated by yellow box in B and showing TFAM localization in red with anti-TFAM); the length of the box = 4 µm. (E-G) Normalized intensity distributions of nucleoids labelled with anti-DNA and with anti-TFAM corresponding to images from E and F, respectively. (H-I) Time-course experiment of live HGPS cells under photo-toxic conditions. Mitochondria were labelled with MitoTracker Deep Red (H, magenta) and mtDNA was labelled with PicoGreen (H,I, green). Scale bar = 2 µm. Arrow heads indicate pairs of nucleoids that undergo liquid-like fusion events. (J-K) Single z-slices of SIM images of UPR^mt^ in mitochondria from fixed normal (J) and an HGPS (K) human skin fibroblast, where the UPR^mt^ marker is gray-scale with anti-HSPD1 (mtHSP60), and mtDNA nucleoids are in green with anti-DNA. Scale bar = 1.5 µm. (O) Immunofluorescence quantification of UPR^mt^ markers (ClpP, LonPI, mtHSP10, mtHSP60, mtHSP70) in all six primary cell lines reported as normalized intensity values in undamaged and damaged mitochondria from high-throughput confocal images, where n=3 independent experimental replicates (each experimental replicate, containing three technical replicates, had a total of 150-600 cells for each cell line and UPR^mt^ marker) and error bars are standard error.

Analysis by high-resolution Structured Illumination Microscopy (SIM) imaging revealed the presence of enlarged nucleoids in HGPS cells (Figures 1F-1G). In morphologically aberrant mitochondria, mt-nucleoids clustered together into structures that were considerably brighter and larger than the typically uniform nucleoids of ∼100 nm found in normal mitochondria (Figures 1D-1E). Several mt-nucleoid markers, including mtDNA and the major mtDNA-packaging protein, TFAM, were locally enriched in the atypical mitochondria (Figures 1F, 1G, S1O and S1P), while total TFAM and mtDNA levels were not altered in HGPS cells (Figures S1L-S1N).

To visualize the formation of enlarged mt-nucleoids *in vivo*, we exposed primary skin fibroblasts to phototoxic stress (Minamikawa et al., 1999) (see Supplementary Information). Within ∼10 minutes, neighboring mt-nucleoids dynamically fused to generate enlarged droplet-like structures greater than 100 nm in size and up to a few microns in diameter analogous to those seen in damaged, swollen mitochondria of HGPS cells (Figures 1H and 1I and Videos S1 and S2). Similar, but even more pronounced, fusion events were observed in the presence of the intercalator EtBr (Videos S3 and S4). The homotypic fusion events between neighboring mt-nucleoids are consistent with the behavior of coalescing liquid droplets (Banani et al., 2017; Hyman et al., 2014). We conclude that loss of mitochondrial homeostasis results in the inability of mitochondria to maintain nucleoid size and leads to the coalescence of multiple proximal mt-nucleoids to form larger droplets of nucleoprotein complexes in a process that closely resembles the phase separation of many other biomolecular condensates (Banani et al., 2017; Hyman et al., 2014).

### TFAM phase separates *in vitro* into viscoelastic droplets

To explore if liquid-liquid phase separation drives mt-nucleoid assembly, we examined the ability of the major nucleoid packaging protein TFAM to undergo phase separation *in vitro*. TFAM phase separated into spherical droplets in low salt conditions and at protein concentrations of ≥5 *μ*M TFAM (Figures 2A and S2D). Droplet formation was reversible upon increasing salt concentration (Figure S2E). After 30-60 min post-mixing, droplets coarsened to sizes of ∼1-5 *μ*m and sedimented towards the bottom of the imaging chamber (Figure 2A). The TFAM concentrations required for phase separation *in vitro* were well within the estimated physiological range inside the mitochondria of ∼10 *μ*M (see Supplementary Information) (Kukat et al., 2011).

**Figure 2:**
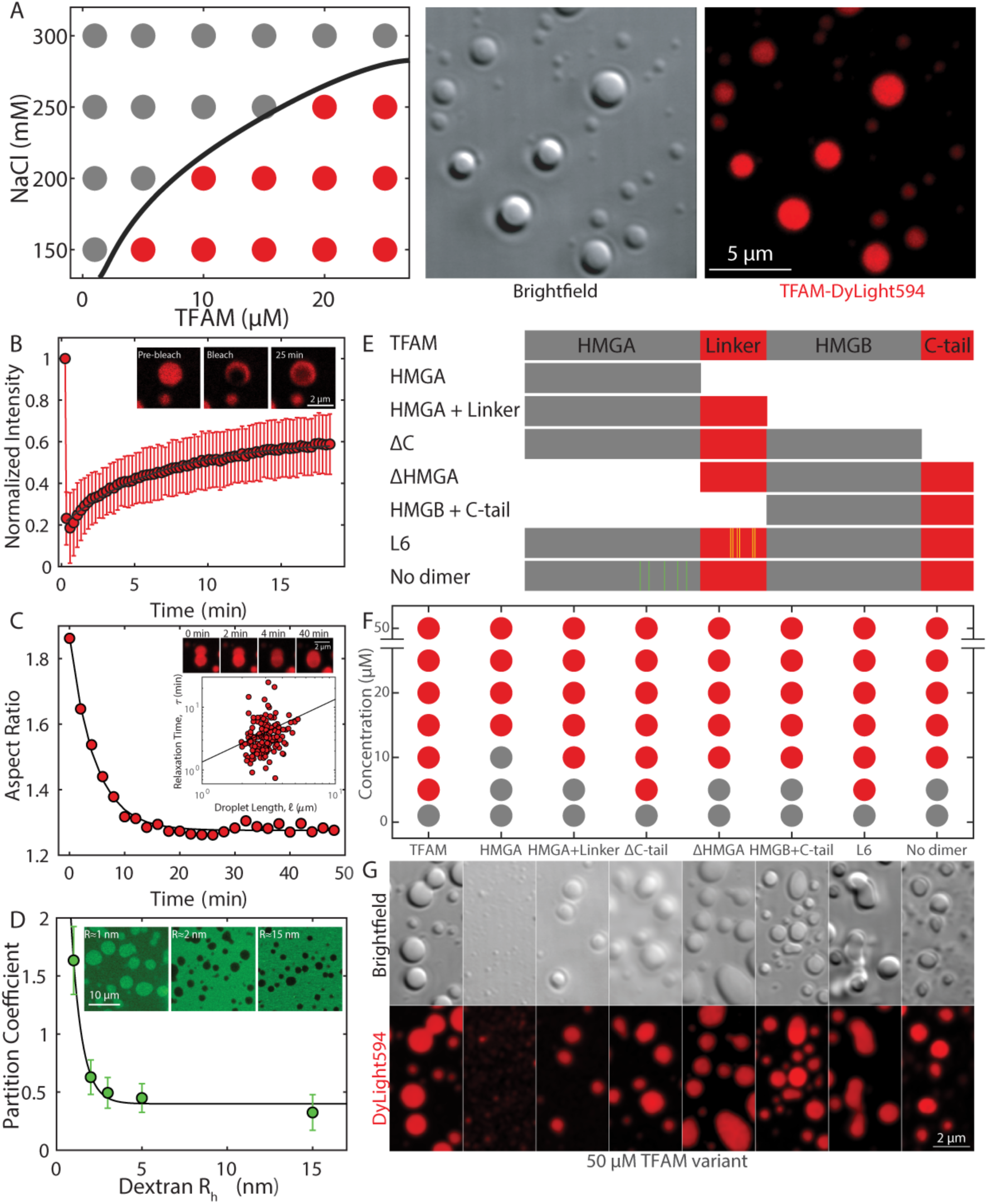
TFAM phase separates into viscoelastic droplets, driven by multivalent interactions. (A) Phase diagram of TFAM under various protein and salt concentrations, where grey dots indicate single/soluble phase, red dots signify two phases/droplets present. DIC image (top) and maximum intensity projection (bottom) of TFAM-DyLight594 droplets at 25 μM and 150 mM NaCl in 20 mM Tris-HCl, pH 7.5 thirty minutes after mixing. Scale bar = 5 μm. (B) FRAP using 488 and 561 nm light performed on a ∼1 μm spot on TFAM droplet thirty minutes after mixing. Inset shows representative fluorescent image of TFAM-DyLight594 pre-bleach, immediately post-bleach, and 25 minutes post-bleach. Scale bar = 2 μm. Values represent averages ± SD from n = 15 droplets. (C) Aspect ratio of droplet shape as a function of time after contact for a representative droplet. Top inset shows fusion images corresponding to the trace at t = 0, 2, 4 and 40 mins. Scale bar = 2 μm. Bottom inset shows data for all droplets analyzed (n= 3 experimental replicates, where ∼50 droplets analyzed per experiment) of characteristic relaxation times as a function of droplet size. Solid line is the linear fit, where the slope is a measure of the inverse capillary velocity. (D) The partition coefficient of dextran-FITC into TFAM droplets as a function of dextran average hydrodynamic radius estimated from the molecular weight. Inset shows representative images showing localization of dextran-FITC for R_*h*_ ≈ 1, 2 and 25 nm. Scale bar = 10 µm. Values represent averages ± SD from n = 3 experiments (>20 droplets analyzed per condition for each experiment). (E) Schematic diagram of mutants with HMG domains in grey and intrinsically disordered regions in red. Yellow and green lines indicate point mutations in L6 and no dimer mutants, respectively. (F) Phase diagram of mutants at 150 mM NaCl and 20 mM Tris-HCl, pH 7.5 for a range of protein concentrations. (G) Fluorescent maximum intensity projections of mutants at 50 μM protein and 150 mM NaCl, 20 mM Tris-HCl, pH 7.5 within 30-60 minutes after mixing. Scale bar = 2 μm.

We performed a series of biophysical assays on TFAM droplets to assess their material properties within the first 30-90 minutes after mixing. Photobleaching of a *R* ≈ 0.5 *μm* spot revealed slow dynamics with a characteristic time scale of *τ* ≈ 6.5 ± 0.5 minutes, which corresponded to a diffusivity of ∼6 × 10^−4^ *μm*^2^/*s*. Furthermore, the immobile fraction of 0.5±0.2 was indicative of viscoelastic behavior (Figure 2B, Video S5). Similarly, time-lapse images of TFAM droplets undergoing coalescence events also displayed slow dynamics with time scales of *τ*=4±0.5 min, giving rise to an inverse capillary velocity of 80±20 s/µm (Figure 2C). Although the droplets had the propensity to relax upon contact, the average aspect ratio upon fusion was 1.36±0.04, which deviated from that of a sphere (AR=1.0) (Figure S2F). Introduction of dextran-FITC of varying sizes as an inert probe to sample the physicochemical environment of the droplets demonstrated that small particles of ≤1 nm preferentially accumulate within the droplets, while increasing probe size resulted in reduced partitioning (Figure 2D). These properties are indicative of a characteristic pore or mesh size of ∼1 nm, suggesting the presence of a polymer meshwork forming amongst individual TFAM molecules within the droplets. These biophysical properties suggest that TFAM molecules form an entangled polymer meshwork or gel with markedly slow internal arrangements and signatures of viscoelasticity.

### TFAM phase separation is driven by multivalent interactions

TFAM contains two DNA-binding High Mobility Group (HMG) domains separated by a disordered linker domain and flanked by an intrinsically disordered C-tail, together forming a relatively flexible chain (Figure S2A) (Ngo et al., 2011; Rubio-Cosials et al., 2011). Additionally, TFAM is one of the most highly charged proteins in the mt-nucleoid (Figure S2C) and is primarily enriched in positive amino acids distributed throughout the length of the protein (Figure S2B). To dissect the molecular features of TFAM responsible for phase separation, a set of TFAM mutants was analyzed for their phase separation behavior *in vitro* (Figure 2E).

In phase separation assays, the HMGA domain alone failed to form the typical micron-sized droplets as seen with full-length TFAM, but assembled into small puncta near the diffraction limit even at high concentrations (Figures 2F, 2G, S2G and S2H). Maintaining half of the protein, either by adding a linker to HMGA (HMGA+linker) or with the analogous HMGB+C-tail mutant, restored droplet formation, albeit at higher saturation concentrations than full length TFAM, suggesting that multi-valency seen in the full-length protein lowers the barrier for phase separation and that the addition of a disordered domain to HMGA promotes phase separation (Figures 2F, 2G, S2G and S2H). Loss of the disordered C-tail (ΔC-tail) did not affect phase separation, but influenced the wetting behavior of the droplets as indicated by a decreased smoothness along the droplet perimeter (Figures 2F, 2G, S2G and S2H), indicating that the disordered C-tail regulates the molecular interactions within the droplet phase. Consistent with this notion, removal of the HMGA domain (ΔHMGA), also resulted in droplet formation, but at slightly higher saturation concentrations. Finally, introduction of six non-polar residues in the linker region (L6 mutant) (Ngo et al., 2014) enhanced the gel-like properties of droplets as evidenced by the highly non-spherical morphologies, whereas inclusion of non-polar residues in the HMGA domain to prevent dimerization (no dimer mutant) (Ngo et al., 2014) increased saturation concentrations and produced smaller droplets, underscoring the contribution of multivalent interactions in phase separation of TFAM (Figures 2F, 2G, S2G and S2H). Taken together, these observations suggest a mechanism by which many weak interactions along a flexible backbone of TFAM allow for robust phase separation and that the disordered linker and C-tail provide flexibility of the biopolymer chain to promote phase separation into prominent droplets.

### Formation of TFAM-mtDNA multiphase, gel-like structures *in vitro*

To examine the interplay of mtDNA and TFAM in phase separated mt-nucleoids, as would occur in the context of mitochondria, we investigated the *in vitro* phase separation behavior of TFAM in the presence of mtDNA (Figures 3, S3A, and S3B). As expected, mtDNA (0-10 nM or equivalently 0-100 ng/ul) on its own did not phase separate, but when combined with TFAM at concentrations that support phase separation (≥5 *μ*M), mtDNA readily partitioned into droplets. Importantly, the presence of mtDNA significantly affected droplet formation and morphology (Figures 3A and S3C). At a DNA/TFAM molar ratio of ∼0.001, which corresponds to estimates of their physiological ratio (Kukat et al., 2011) (see Supplementary Information), TFAM and mtDNA readily formed droplets (Figures 3A and S3C). At high mtDNA/TFAM molar ratios of >0.001, droplets ceased to form, potentially due to saturation behavior (Figures 3A and S3C). For molar ratios of mtDNA/TFAM ≤6E-4, the aspect ratio of the droplets notably increased with increasing ratio of DNA/TFAM mass concentrations (Figure 3B). For mtDNA/TFAM molar ratios <3E-4, the number of droplets (measured ∼1 hour after mixing) increased with increasing mtDNA/TFAM levels (Figure 3B inset), suggesting that mtDNA can potentiate droplet formation under those conditions, possibly acting as a nucleating agent and paralleling how RNA drives phase separation when added to RNA-binding proteins (Lin et al., 2015). ssDNA, dsDNA, and RNA as well as free nucleotides (dNTPs) also supported TFAM droplet formation (Figures S3D-S3S). Interestingly, ssDNA and dsDNA, considerably longer than the 16.6 kB mtDNA and with no sequence specificity, resulted in even more irregular droplet morphologies and pronounced gelation (Figures S3D-S3S). We conclude that the addition of long, polymerized strands of DNA, irrespective of sequence, leads to favorable interactions between TFAM and DNA, thereby affecting the emergent droplet behavior (Figures S3D-S3U). These findings demonstrate that the material properties of the droplets depend on DNA/TFAM composition, where increasing DNA promotes gelation.

**Figure 3:**
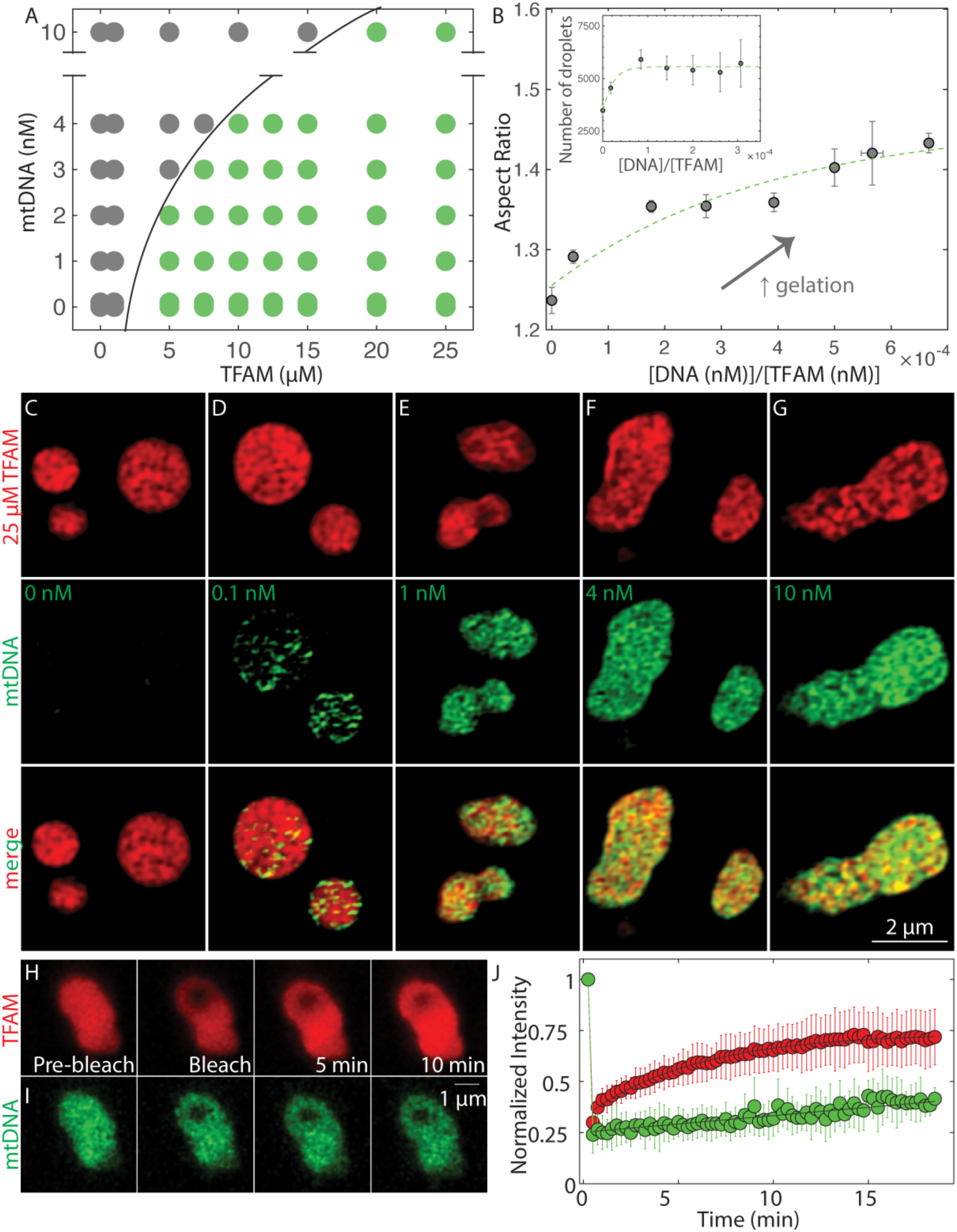
Formation of TFAM-mtDNA multiphase, gel-like structures. (A) Phase diagram of mtDNA versus TFAM denoting single/soluble phase (gray) or two phases/droplets (green). Each point on the phase diagram representing a unique DNA and protein concentration was measured from n=2-12 independent experiments. Black solid line delineates deduced phase boundary. (B) Aspect ratio as a function of dimensionless concentration (molar concentration DNA/molar concentration TFAM). Values represent binned conditions from DNA/TFAM conditions measured in (A) and error bars are SEM. Inset: number of TFAM-mtDNA droplets per field of view as a function of dimensionless concentration. Values represent binned conditions from DNA/TFAM conditions measured in (A) and error bars are SEM. (C-G) SIM images of droplets thirty minutes after mixing with various amounts of mtDNA: 0 nM (C), 0.1 nM (D), 1 nM (E), 4 nM (F), and 10 nM (G). Top row is of TFAM-DyLight594 (red), middle row is of mtDNA-Alexa488 (green) and bottom row is the merged image. Scale bar = 2 μm. (H-I) FRAP experiments on TFAM-mtDNA droplets at 25 µM TFAM (H, red) and 10 nM mtDNA (I, green). (J) FRAP recovery curve showing intensity as a function of time for TFAM (red) and mtDNA (green). Values represent averages ± SD from n = 16 droplets.

To probe how mtDNA localizes within the droplets, we performed SIM imaging on TFAM-mtDNA droplets containing increasingly higher concentrations of mtDNA. We find that mtDNA is not uniformly distributed, but de-mixes from the majority of TFAM within the droplet (Figures 3C-3G), consistent with multiphase behavior seen in other multi-component phase separating systems, such as the nucleolus (Feric et al., 2016). Multiphase organization was observed with both ssDNA and dsDNA, but not with dNTPs nor RNA (Figures S3J-S3S). To characterize the dynamics of multiphase TFAM-mtDNA droplets, fluorescence recovery after photobleaching (FRAP) demonstrated that TFAM was able to diffuse within TFAM-mtDNA droplets with similar recovery behavior as in pure TFAM droplets (Figures 3H and 3J, Video S6). In contrast, on these timescales, mtDNA within the droplets remained strikingly immobile (Figures 3I and 3J, Video S6), suggesting that the mtDNA molecules within the TFAM-mtDNA droplets determine the time scale for relaxation of the droplets, while also explaining the observed non-spherical shapes at high mtDNA/TFAM ratios. The observed dynamics are consistent with the multiphase sub-structure of these droplets.

### Phase behavior of TFAM in live cells

To test if the *in vitro* phase-separation behavior of TFAM and mtDNA reflects the dynamic properties and structural organization of nucleoids *in vivo*, we visualized mt-nucleoids in HeLa cells using TFAM-mKate2. In photobleaching experiments, TFAM-mKate2 exhibited very low recovery (immobile fraction

= 0.9±0.3) (Figures S4A-D, Video S7) indicative of limited exchange of TFAM between the mt-nucleoid and the mitochondrial volume, consistent with the very low concentration of free TFAM (Lu et al., 2013; Matsushima et al., 2010). However, in line with *in vitro* observations, the ability of TFAM to rapidly diffuse within a nucleoid became evident when a bleached nucleoid fused with a neighboring unbleached nucleoid resulting in rapid exchange within the coalescing droplet (Figures 4A and 4B, Video S8).

**Figure 4:**
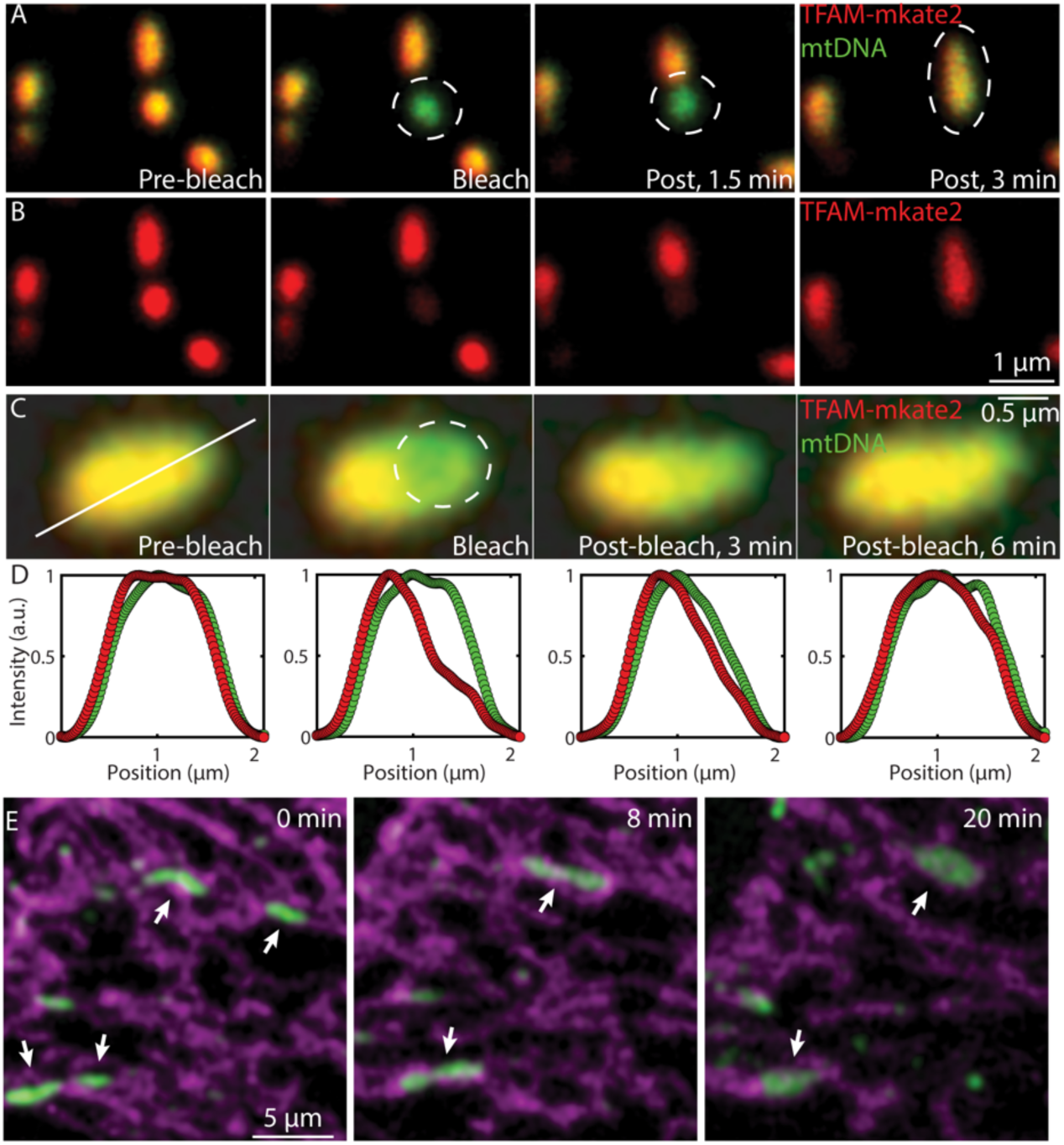
Phase separation behavior of TFAM in live cells. (A-B) FRAP experiment performed on a small nucleoid that later undergoes a fusion event in live HeLa cells expressing TFAM-mKate2. (A) The overlay of TFAM-mKate2 (red) and of mtDNA labelled with PicoGreen (green) pre-bleach, bleach, 1.5 minutes and 3 minutes post-bleach, and (B), the single-channel of TFAM-mKate2 (red). Scale bar = 1 µm. (C-D) FRAP experiment performed on an enlarged nucleoid in live HeLa cells expressing TFAM-mKate2 (n = 18 cells), where (C) is the overlay of TFAM-mKate2 (red) and of mtDNA labelled with PicoGreen (green) pre-bleach, bleach, 3 minutes and 6 minutes post-bleach. The solid white line indicates the pixels that were analyzed for intensity quantification. The dashed white circle denotes the part of the nucleoid that was bleached. Scale bar = 0.5 µm. (D) The normalized intensity of TFAM-mKate2 (red) and PicoGreen (green) where the x-axis corresponds to the solid white line from (C). (E) Images of mt-nucleoids undergoing liquid-like fusion events in HeLa cells overexpressing TFAM-mKate2 after ∼30 minute exposure to EtBr with phototoxic imaging conditions where mtDNA (PicoGreen, green) and mitochondria (MitoTracker Deep Red, magenta) are shown. Arrow heads point to nucleoids involved in homotypic fusion events. Scale bar = 5 µm.

Furthermore, overexpression of TFAM-mKate2 to high levels in HeLa cells led to formation of enlarged nucleoids (∼1 *μm*) (Figures S4F and S4G), and photobleaching a small spot (*R* ≈ 0.2 μm) within these structures (Figures 4C and S4E, Video S9) gave rise to recovery of ∼70±20% of signal further indicating relatively high mobility of TFAM within mt-nucleoids (Figure 4D). Based on observed recovery times of ∼1 min, we estimate diffusivities of ∼1 × 10^−3^ *μm*^2^/*s* (Figure 4D), in agreement with *in vitro* observations. Moreover, nucleoids larger than the diffraction limit exhibit clear liquid-like fusion behavior upon phototoxic and EtBr induced stress, consistent with a phase separation model (Figure 4E, Video S10), and exist as discrete structures in sporadically swollen mitochondria in HeLa cells (Figure S4J). Furthermore, mt-nucleoids were largely unaffected upon swelling of mitochondria under hypo-osmotic conditions (Figures S4H and S4I), suggesting nucleoid components are not diffuse in the mitochondrial matrix, but that the size and shape of mt-nucleoids are intrinsic properties arising from phase separation. These dynamics and recovery features are comparable to the viscoelastic material properties of TFAM droplets *in vitro*.

### TFAM influences localization and miscibility of mt-nucleoid components

Our *in vitro* titration experiments suggested correlation between TFAM and mtDNA concentrations and morphology of the phase separated structures. To probe the effect of TFAM concentration on mt-nucleoid organization *in vivo*, we expressed TFAM-mKate2 and selected nucleoids with varying levels of expression for analysis using SIM. Consistent with *in vitro* titration data, we find an increase of nucleoid sizes, reaching up to a few microns in length, upon increasing levels of TFAM-mKate2 (Figures 5A-5C and S5A). In addition, the mtDNA localization within these enlarged nucleoids had similar multiphase organization as observed in TFAM-mtDNA droplets *in vitro* (Figures 5A-5C). This non-uniform structure is consistent with the layered organization deduced from biochemical analysis (Bogenhagen et al., 2008).

**Figure 5:**
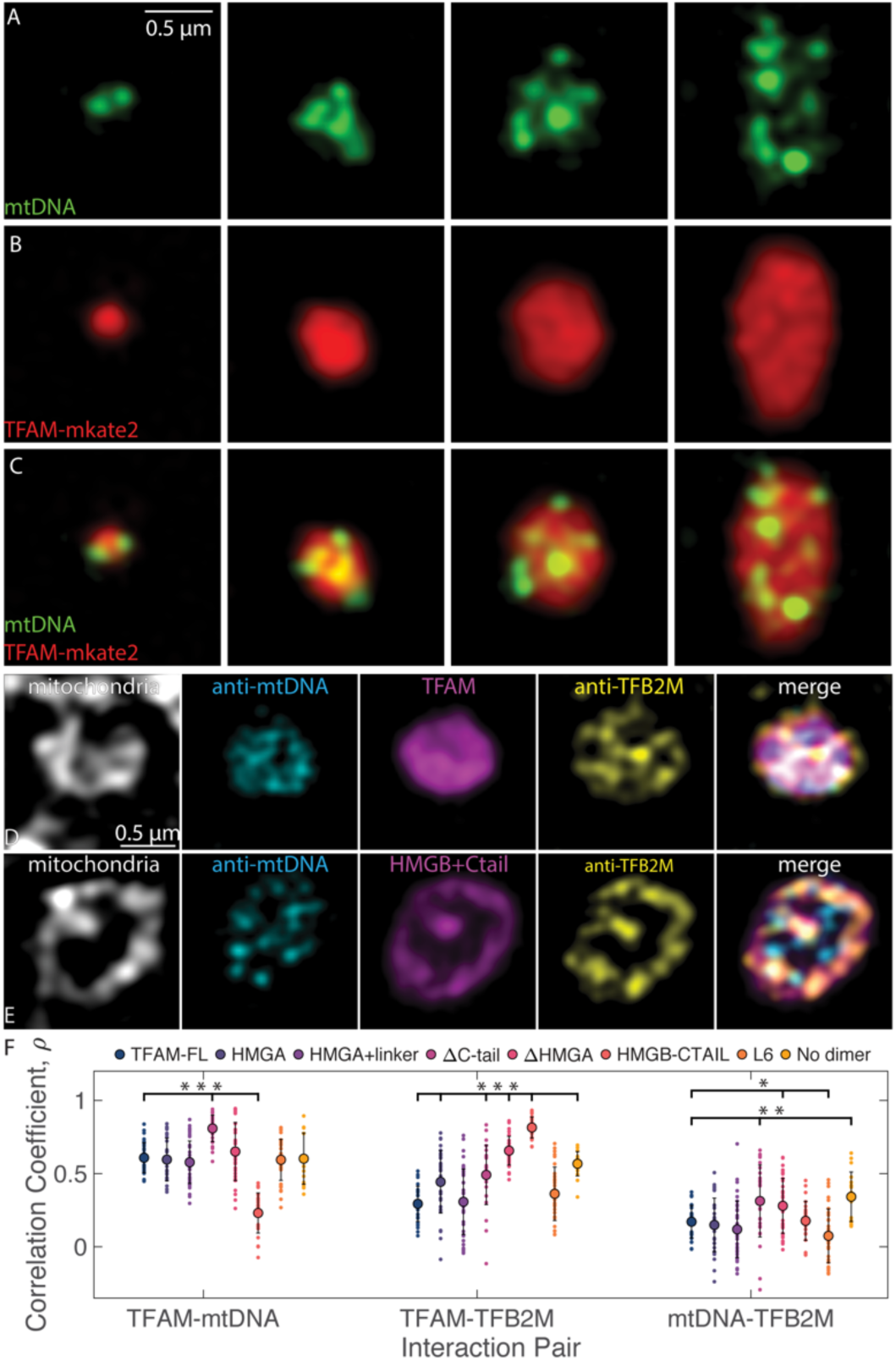
TFAM mutants influence miscibility and localization of mt-nucleoid components. (A-D) Single z-slices of SIM merged images of nucleoids ranging in size after overexpression of TFAM-mKate2 in a fixed HeLa cell, where mtDNA (A, green, anti-DNA), and TFAM (B, red, TFAM-mKate2) and merge (C). Scale bar = 0.5 µm. (D-E) Single z-slices of SIM individual channels of fixed HeLa cells after TFAM-mKate2 (D) and HMGB+C-tail-mKate2 (E) overexpression (magenta), where mitochondria are labelled with MitoTracker Far Red (gray), mtDNA is labelled with anti-mtDNA (cyan) and TFB2M is labelled with anti-TFB2M (yellow). Merged images are shown as an overlay of cyan, magenta, and yellow channels only. Scale bar = 0.5 µm. (F) Colocalization of channels is shown computed from Pearson’s correlation coefficient for each interaction pair shown for TFAM-FL and all TFAM mutant constructs overexpressed in HeLa cells (n=20-40 nucleoids from 4-5 cells imaged for each mutant construct). ANOVA analysis shows statistical significance across all groups, where p<2e-16 for TFAM-mtDNA interactions, p<2e-16 for TFAM-TFB2M interactions, and p = 1.3e-8 for mtDNA-TFB2M interactions. Notation on graph denotes statistical analysis relative to TFAM-FL, where *p<0.05, **p<0.01, and ***p<0.001.

We next characterized the effect of TFAM on the multi-phase properties of mt-nucleoids by analyzing the behavior of an additional core mitochondrial transcription factor, TFB2M (Bogenhagen et al., 2008). We find that TFB2M did not mix homogenously with TFAM and DNA, but was preferentially localized towards the mitochondrial inner membrane (Figure 5D) in line with the multi-component and multi-phase nature of mt-nucleoids. Based on Pearson’s correlation coefficient to assess partitioning within each microphase, TFAM has higher colocalization with mtDNA (*ρ*_*TFAM,mtDNA*_=0.6±0.1) than with TFB2M (*ρ*_*TFAM,TFB2M*_ =0.3±0.1) (Figure 5F). Furthermore, the correlation between TFAM and mtDNA is not 1, suggesting that there are microdomains that are enriched in TFAM, consistent with the *in vitro* multiphase behavior.

To relate the *in vitro* behavior of mutant TFAM to its *in vivo* properties, we examined a series of TFAM mutants in HeLa cells (Figures 5E and S5B-S5G). Notably, the HMGB+C-tail-mutant caused a complete inversion in localization of the mutant from the interior into the lining of the membrane (*ρ*_*HMGB+C-tail,mtDNA*_=0.2±0.1, *ρ*_*HMGB+C-tail,TFB2M*_=0.8±0.1) (Figure 5F). This change in localization reflected differences in protein miscibility of HMGB+C-tail between the microphases, demonstrating that the HMGA and linker domains are important for association of TFAM with mtDNA. Other mutants also affected interactions between the mt-nucleoid components, although to a lesser extent. Specifically, the ΔC-tail mutant increased colocalization with mtDNA (*ρΔC-*_*tail,mtDNA*_ =0.8±0.1), which suggests that loss of the C-tail increased affinity and binding of TFAM and mtDNA, while potentially competing for interactions with other proteins (Figures 5F and S5D). Several other mutants also had increased affinities for TFB2M, albeit to a lesser extent as the HMGB+C-tail mutant, including the ΔHMGA mutant (ρ_***ΔHMGA***,***TFB2M***_=0.7±0.1), which approached HMGB+C-tail in composition (Figures 5F and S5E).

Taken together, these results demonstrate that the *in vitro* properties of TFAM-mtDNA multiphase structures reflect the underlying physics of mitochondrial nucleoids *in vivo*.

### Functional features of enlarged mt-nucleoids

To finally assess whether the phase-separation properties of mt-nucleoids are related to mitochondrial function, we measured mitochondrial activities in HGPS cells, which are enriched for enlarged mt-nucleoids (Figure 6). Single molecule FISH for mt-12S and mt-COI RNA demonstrated enrichment of mt-RNA transcripts in enlarged nucleoids that were proportional to local TFAM levels (Figures 6A, 6B, S6A-S6F, and S6M). RNA transcripts in enlarged mt-nucleoids localized along the perimeter of the mitochondrial membrane, but did not colocalize with nucleoids, suggesting that nucleoids and mt-RNA exist as distinct structures, and potentially immiscible phases, of (ribo)nucleoproteins (Figures 6C-6F), which is in line with the weak partitioning and minor effect on morphology of RNA relative to DNA on TFAM droplets in the *in vitro* system (Figures S3D-S3U). Similar increased transcriptional activity of enlarged mt-nucleoids was evident when nascent transcription was measured by BrU incorporation (Figures S6G-S6L and S6N). Consistently, we did not observe changes in mt-RNA transcripts on a population level as measured by qPCR, which supports that enlarged nucleoids remain transcriptionally active, also in line with RNA-FISH imaging (Figure S6O). However, the enrichment of enlarged phase-separated mt-nucleoids in HGPS was associated with altered mitochondrial metabolic functions (Figure 6G). Basal mitochondrial respiration, maximal respiration and reserve capacity were reduced in HGPS fibroblasts containing enlarged mt-nucleoids compared to isogenic non-affected fibroblasts (Figures 6H-6J), indicating functional impairment of mitochondrial oxidative phosphorylation and ATP regeneration. Furthermore, HGPS fibroblasts from older affected donors had elevated mitochondrial ROS and membrane potential compared to young HGPS or unaffected proband control fibroblasts (Figures 6K and 6L). These observations demonstrate that the presence of enlarged mt-nucleoids generated by phase separation is accompanied by mitochondrial dysfunction.

**Figure 6:**
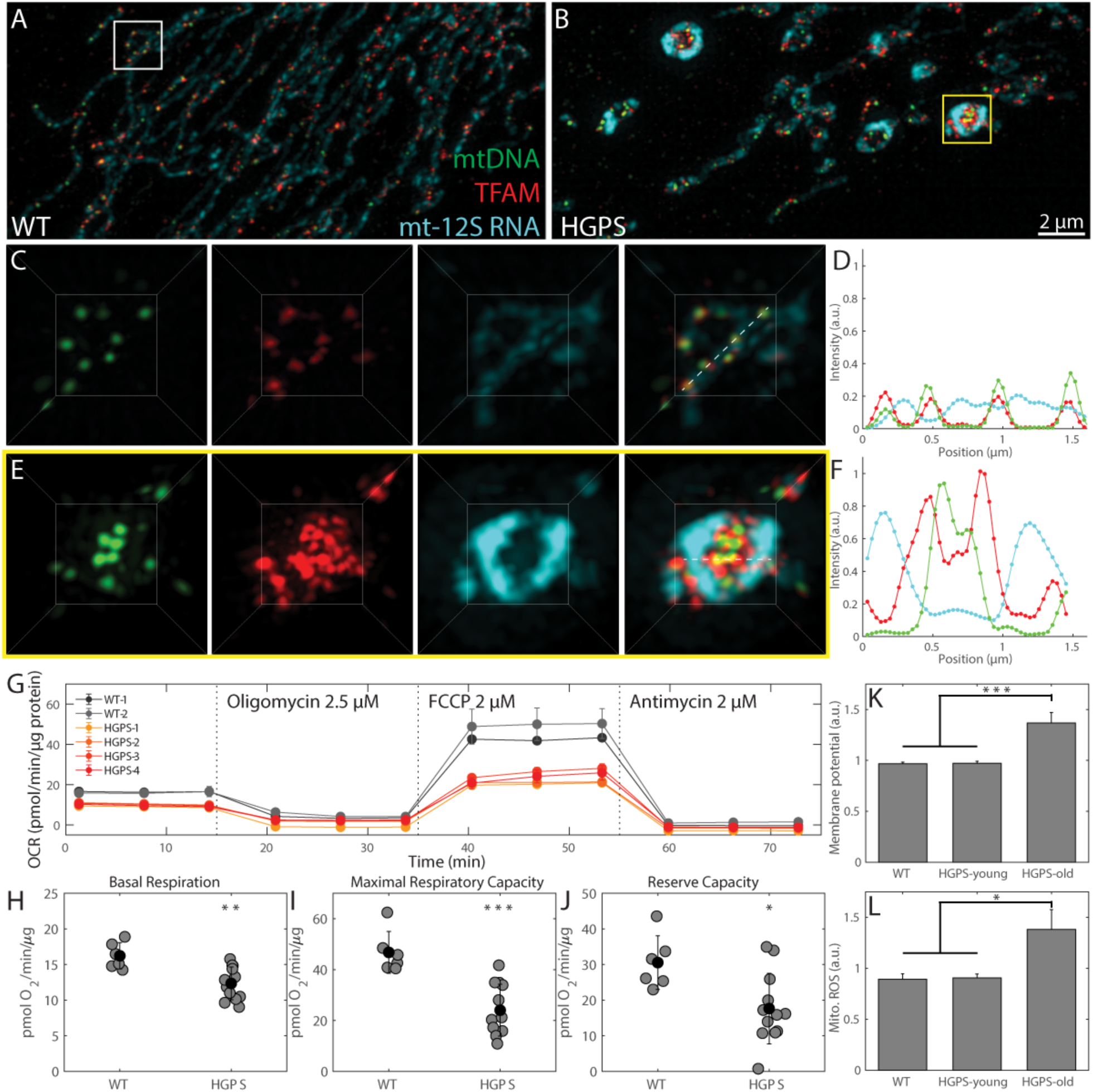
Functional features of enlarged mt-nucleoids. (A-B) Maximum intensity projections of SIM images of a fixed normal (A) and an HGPS (B) human skin fibroblast, where mtDNA is in green with anti-DNA, TFAM is in red with anti-TFAM, and mt-12S RNA FISH is in cyan. Scale bar = 2 µm. (C,E), Three-dimensional views of normal mitochondria (C) annotated by white box in A and swollen mitochondria (E) annotated by yellow box in (B) and showing individual channels and an overlay; box ≈ 2.5 x 3 µm. (D,F) Normalized intensity distributions of nucleoids labelled with anti-DNA (green), anti-TFAM (red), and mt-12S RNA (cyan) corresponding to images from (C) and (E), respectively. (G-J) Seahorse assay results on primary skin fibroblasts from wildtype cell lines (WT-1,2) and HGPS cell lines (HGPS-1,2,3,4) from a representative experiment. (G) The oxygen consumption rate (OCR) as a function of time after perturbation with oligomycin, 2.5 µM at t = 16 minutes, FCCP at 2 μM, at t = 24 minutes, and antimycin 2 μM at t = 55 minutes. Representative trace from a single experiment, error bars are SEM of technical replicates (n=3). Averaged results for WT and HGPS cells pooled together: basal respiration (H), maximal respiratory capacity (I), and reserve capacity (J). Error bars are standard deviation, where *p<0.05, **p<0.01 and ***p<0.001. For (G), three experimental replicates were performed, and for (H-J), experiments were pooled among cell types, where values represent averages ± SD from n = 6 independent experimental replicates of WT cells and n = 12 independent experimental replicates of HGPS cells. (K) Mitochondrial membrane potential using TMRM. Cells were grouped as WT (n=8 independent measurements), HGPS young (n=8 independent measurements), and HGPS old (n=8 independent measurements). Error bars are SEM, where p-value for the ANOVA test statistic was p<0.001. For individual pairs, ***p<0.001. (L) Mitochondrial ROS using MitoSOX Red. Cells were grouped as WT (n=6 independent measurements), HGPS young (n=6 independent measurements), and HGPS old (n=6 independent measurements). Error bars are SEM, where p-value for the ANOVA test statistic was p<0.05. For individual pairs, *p<0.05.

## Discussion

Our results show that mitochondrial nucleoids form by phase separation. Our finding that the well-established architectural mt-nucleoid protein TFAM has a propensity for self-assembly into a separate protein-rich phase and, combined with mt-DNA, generates multi-phasic structures *in vitro* suggests that TFAM is the major driver of mt-nucleoid phase separation. In support of this, we find that the properties of *in vitro* generated condensates containing TFAM and mtDNA mimic *in vivo* behavior of mt-nucleoids. These results provide new insights into the biogenesis of mt-nucleoids and the function of the major architectural mtDNA-binding protein TFAM. While the ability of TFAM to bind DNA is well established (Farge et al., 2014; Kukat et al., 2015; Wong et al., 2009), binding of TFAM to mtDNA alone does not explain the higher-order structural features of mt-nucleoids, such as their observed size and shape, and their dynamic emergent properties, including liquid-like fusion events and internal rearrangements. The phase behavior of TFAM and mtDNA described here accounts for the observed morphological features of mt-nucleoids as discrete, heterogenous, non-membranous entities within mitochondria, their viscoelastic dynamics *in vivo*, and their ability to reach sizes larger than 100 nm as observed in HGPS cells. This physical phase separation model complements previous self-assembly studies of TFAM and mtDNA into nucleoid-like structures under dilute conditions (Farge et al., 2014; Kukat et al., 2015; Wong et al., 2009), by extending them to reveal the importance of protein-protein interactions necessary in driving the self-assembly of the mitochondrial genome.

Based on our mutational analysis of TFAM, we suggest a model in which weak interactions along the flexible backbone of TFAM promote phase separation into viscoelastic droplets. In the presence of DNA, the HMG domains of TFAM each bind DNA by intercalating into the helical strand, and together, further bend and stabilize the DNA fiber (Figure S4B) (Ngo et al., 2011; Rubio-Cosials et al., 2011).

Collectively, TFAM molecules fully coat the DNA fiber, encapsulating it within a gel meshwork, while interacting with other bound or unbound TFAM molecules to form a phase separated condensate. Differential interactions and solubilities of full length TFAM with nucleoid components (mtDNA, TFB2M) likely contribute to the generation of microphases within the mt-nucleoid droplet and the multi-phasic nature of mt-nucleoids. Our observation that truncation and modification of TFAM domains leads to pronounced changes in localization within the nucleoid droplet underscores the role of TFAM-specific interactions in independently conferring miscibility and determining nucleoid structure. These physical properties may also generate a platform for other mt-nucleoid associated proteins to partition into these dynamic, yet persistent, structures (Bogenhagen et al., 2008) and to exclude other components from the mitochondrial matrix from fully mixing with the mt-nucleoid.

The observed preferential localization of a TFAM mutant consisting of the HMGB- and C-terminal domain with the mitochondrial membrane suggests that the TFAM N-terminus interacts with mtDNA in the nucleoid interior within the mitochondrial matrix, and that the C-terminus is relatively less soluble and interacts with the mitochondrial inner membrane. Previous work suggested that mitochondrial nucleoids interact with membranes (Brown et al., 2011; Kopek et al., 2012), potentially via a tethering mechanism (Chen and Butow, 2005). Our results suggest that the interaction of nucleoids with membranes is due to the emergent wetting behavior of the condensate on the membrane (Snead and Gladfelter, 2019). Similar observations have been made for contact sites between tethering proteins from various membrane bound organelles, including the mitochondrial mitofusin 1 (Mfn1) tethering protein and Sec61β of the ER membrane (King et al., 2020), and for the condensation of Atg1-complex droplets, as part of the pre-autophagosomal structure (PAS), along vacuolar membranes (Fujioka et al., 2020). It is tempting to speculate that the wetting behavior between the mitochondrial inner membrane and the nucleoid may not only play a role in regulating the size and diffusion of nucleoids, but also in functional processes such as replication (Lewis et al., 2016). In addition, mitochondrial nucleoids may not be the only example of phase separation within the mitochondria, as mitochondria have also been reported to contain various RNA granules (Jourdain et al., 2016; Jourdain et al., 2013), which do not colocalize with mt-nucleoids. This is further supported by our observation that RNA weakly partitions into TFAM droplets *in vitro*, suggesting poor miscibility of RNA within the mt-nucleoid phase.

We find that enlarged mt-nucleoids in HGPS cells are associated with mitochondrial dysfunction. Our observation that transcriptional levels are unchanged in enlarged nucleoids is consistent with phase separation: if both small and large droplets arise from the same coordinate on the phase diagram, then the droplets should be compositionally and functionally similar. We find enlarged nucleoids under conditions of acute or chronic stress such as phototoxicity or as in HGPS cells, which exhibit elevated levels of ROS (Kubben et al., 2016). An intriguing possibility with regards to functional properties of mt-nucleoids is that phase separation of TFAM may be a protective response towards maintaining mtDNA sequence integrity (Bogenhagen, 2012; Cadenas and Davies, 2000). The formation of a gel-like proteinaceous layer of TFAM around mtDNA may act as a physical barrier impeding diffusion of free radicals. Fusion into a single larger, spherical droplet would reduce the total surface area exposed to the surrounding oxidizing environment and increase the distance for ROS to diffuse to reach the mt-nucleoid core, thereby minimizing the formation of ROS-induced mutations and deterioration of the mitochondrial genome (Barja and Herrero, 2000). Additionally, phase separation of the mt-nucleoid components into droplet-like structures could serve as a platform to recruit and retain necessary mtDNA repair factors in response to damage (Kazak et al., 2012).

The phase separation behavior of mt-nucleoids may also account for the observation of a high degree of pathogenicity of a remarkably diverse set of mutations in mt-nucleoid proteins (Suomalainen and Battersby, 2018). For example, a large number point mutations in the linker region of POLG lead to a spectrum of diseases (Chan and Copeland, 2009). These mutations may exert their pathogenic effect not by specifically disrupting the active site of the affected protein, but rather by globally influencing the biophysical interactions that contribute to higher-order structure and therefore long-term stability of mt-nucleoids.

Mitochondria are originally derived from prokaryotes, and nucleoids are an evolutionarily ancient feature used to organize both prokaryotic and mitochondrial genomes (Dillon and Dorman, 2010). In support of a common organizational principle, bacterial genomes are packaged into nucleoid structures that have also been described to behave as fluids (Cunha et al., 2001), and some bacterial nucleoprotein complexes also undergo liquid-liquid phase separation (Monterroso et al., 2019). Unlike TFAM, bacterial architectural nucleoid-associated proteins, including HU, histone-like nucleoid structuring protein (H-NS), factor for inversion stimulation (FIS), and the integration host factor (IHF), lack HMG domains as seen in TFAM, suggesting a multitude of molecular interactions can condense DNA (Dillon and Dorman, 2010; Kucej and Butow, 2007). Interestingly, HMG domains have purely eukaryotic origins, which is also exemplified by how mitochondrial nucleoids in budding yeast are analogously packaged primarily by the TFAM-homologue Abf2p, which also contains two HMG domains separated by a shorter linker (Brewer et al., 2003).

Phase separation has recently been implicated in the organization on the nuclear genome (Gibson et al., 2019; Larson et al., 2017; Strom et al., 2017). DNA in the eukaryotic nuclear genome is wrapped by histone octamers to form nucleosomes that collectively comprise the chromatin fiber. Purification of nucleosomes *in vitro* leads to liquid droplet formation via phase separation and droplet dynamics and organization can be directly modulated by regulatory factors, including linker length and post translational modifications (Gibson et al., 2019). *In vivo*, many chromatin domains are emerging to behave as molecular condensates, including transcriptionally relevant super-enhancers (Sabari et al., 2018). In fact, RNA polymerase II and many nuclear transcription factors have low-complexity domains or intrinsically disordered regions, similar to the disordered modular domains of TFAM and which are features known to promote phase separation (Boehning et al., 2018; Boija et al., 2018; Cho et al., 2018; Chong et al., 2018). On an even larger scale, phase separation can explain the emergence and maintenance of heterochromatin, as HP1a readily phase separates *in vitro* and has dynamic, liquid-like properties in live cells (Larson et al., 2017; Sanulli et al., 2019; Strom et al., 2017). The involvement of phase separation in the organization of diverse genomes from simple mitochondrial and bacterial nucleoids to complex eukaryotic nuclear genomes suggest phase separation is an evolutionarily conserved mechanism for genome organization.

## Acknowledgements

We thank the members of the Misteli lab for discussion and experimental design; L. Schiltz and A. Schibler for initial help with protein purification; J. Jones and M. Taylor for help with final protein expression and purification at the NIH/NCI/CCR Protein Production Core; T. Karpova and D. Ball for help with Structured Illumination Microscopy and Laser Scanning Confocal Microscopy as part of the NIH/NCI/CCR LRBGE Optical Imaging Core; G. Pegoraro and L. Ozbun for help with high-throughput imaging and automated liquid handling at the NIH/NCI/CCR High Throughput Imaging Facility (HiTIF); and K. M. McKinnon for help with cell sorting as part of the NIH/NCI/CCR FACS Core Facility.

## Author contributions

MF performed live/fixed cell microscopy and *in vitro* experiments and analysis. TD, JT, and DC performed and analyzed mitochondrial Seahorse assays, mitochondrial membrane potential and mitochondrial ROS measurements on normal and HGPS cells. TM and MF planned experiments, discussed results, and wrote the manuscript. All authors revised the manuscript.

## Supplementary Information

Methods

References

Supplementary Table S1

Supplementary Figures S1-S6

Videos S1-S10

