## Supplemental Information for "Self-assembly of multi-component mitochondrial nucleoids via phase separation"

**Supplementary Information**

This PDF file includes:

Methods
References
Supplementary Table S1
Supplementary Figures S1 to S6
Captions for Videos S1 to S10

Other Supplementary Materials for this manuscript include:

Videos S1 to S10

### Materials and Methods

#### Cell culture

Primary human dermal fibroblast cell lines were obtained from the Progeria Research Foundation (PRF). Two of the cell lines were from normal, middle-aged parents. The remaining four cell lines were from patients that have the classic mutation in *LMNA* Exon 11, heterozygous c.1824C>T (p.Gly608Gly) and the age at donation sampled early (age 2 and 3 years) and late (ages 6 and 8 years) stages of the disease. For all experiments, cells were passage matched between P10-20. Cells were grown according to PRF's recommended protocols. Briefly, cells were grown in media containing DMEM (Thermo Fisher Scientific, #11960-044) and supplemented with 15% FBS (Thermo Fisher Scientific, #10437), 1% penicillin-streptomycin (Thermo Fisher Scientific), and 1% glutaMAX (Thermo Fisher Scientific, #35050-061) in T25 and T75 mL flasks, and cells were periodically split with trypsin (Thermo Fisher Scientific).

For live cell experiments and for pharmacological disruption experiments, HeLa cells (ATCC) were used. Cells were grown in media containing DMEM (Thermo Fisher Scientific, #11960-044) and supplemented with 10% FBS (Thermo Fisher Scientific, #10437), 1% penicillin-streptomycin (Thermo Fisher Scientific), and 1% glutamine (Thermo Fisher Scientific).

For viral production, HEK293FT cells were used and grown in media containing DMEM (Thermo Fisher Scientific, #11960-044) and supplemented with 10% FBS (Thermo Fisher Scientific, #10437), 1% penicillin-streptomycin (Thermo Fisher Scientific) and 1% glutamine (Thermo Fisher Scientific).

For isolation of mtDNA, immortalized wildtype human fibroblasts CRL-1474 cells were used and grown in media containing MEM (Thermo Fisher Scientific) and supplemented with 15% FBS (as purchased), 1% penicillin-streptomycin (Thermo Fisher Scientific), 1% glutamine, 1% non-essential amino acids (Thermo Fisher Scientific), and 1% sodium pyruvate (Thermo Fisher Scientific). For overexpression of GFP-LA and GFP-Δ50 experiments, immortalized wildtype human fibroblasts CRL-1474 cells were used and grown in media containing DMEM and supplemented with 15% FBS, 1% penicillin-streptomycin, and 1% glutaMAX.

**Lentiviral construct and transfection**

The sequences for TFAM and mKate2 were sequentially cloned from a previous construct (pLenti-GIII-CMV-RFP-2A-Puro, ABM) into a pCDH-CMV backbone (Addgene plasmid #72265) to create the fusion TFAM-mKate2 construct for lentiviral transfection. TFAM mutants (see below, HMGA (43-122), HMGA+linker (43-152), ΔC (43-222), ΔHMGA (123-246), and HMGB+C-tail (153-246), L6 and no dimer) were cloned into the pCDH-CMV-mKate2 vector containing the original TFAM mitochondrial targeting sequence and cleavage site. GFP-Lamin A (GFP-LA) and GFP-Progerin (GFP-Δ50) constructs were in a pCDHblast MCSNard vector, and were a gift from Nard Kubben(DeBoy et al., 2017; Kubben et al., 2012).

HEK293 cells were transfected with vectors containing the gene of interest, pMD2g, and pSPAX2 using PEI. Viruses were collected and concentrated with Lenti-X Concentrator (Takara), flash frozen in liquid nitrogen, and stored in -80C. Frozen aliquots of lentiviruses generated from HEK293 cells were quickly thawed in a 37C water bath, and lentiviruses were applied to HeLa cells and removed after 24 hours. CRL1474 cells were infected with GFP-LA and GFP-Δ50 viruses for 24 hours and were sorted based on GFP expression using a standard FACS protocol. Sorted cells were imaged 2-4 weeks after initial infection.

**Protein constructs and purification**

Human_TFAM_NoMTS_pET28 (full length TFAM protein), Human_TFAM_noMTS_L6_pET28a+ (L6 mutant), and Human _TFAM_NoMTS_Dimer_pET28a+ (no dimer mutant) (Addgene plasmids #34705, #60012, and #60013, respectively) were used. The L6 construct contains the following mutations K136A, H137A, K139A, R140A, K146A, and K147A in the linker region (Ngo et al., 2014). The no-dimer construct contains the following mutations K95A, Y99F, E106A, E112A, and R116A in the HMGA domain to prevent dimerization (Ngo et al., 2014). All other domain mutants were cloned into a pET28a+

expression vector (Novagen) using BamHI and XhoI sites using standard molecular cloning techniques. Specifically, domain mutants consisted of HMGA (43-122), HMGA+linker (43-152), ΔC (43-222), ΔHMGA (123-246), and HMGB+C-tail (153-246) (Ngo et al., 2011).

All proteins were purified using a slightly modified protocol (Ngo et al., 2011). Briefly, all constructs were transformed in BL21 star (DE3) pRare *E. coli*. Bacterial cultures (500 ml for full length TFAM and 250 ml for each mutant) of Dynamite media plus kanamycin and chloramphenicol were inoculated with each construct and incubated until an OD of 7 was reached. IPTG was added at 0.5 mM to induce protein expression, and the culture was incubated overnight at 16C at 220 rpm. The culture was centrifuged and the bacterial pellet was resuspended and lysed in lysis buffer [20 mM Tris-HCl, 500 mM NaCl, pH 8.0]. Cells were mechanically lysed with a microfluidizer, and centrifuged for 30 minutes at 70,000 g at 4C. Protein was purified using an immobilized metal affinity chromatography (IMAC) column, and eluted with elution buffer [20 mM Tris-HCl, 500 mM NaCl, 250 mM Imidazole, pH 8.0], and dialyzed into lysis buffer [20 mM Tris-HCl, 500 mM NaCl, pH 8.0] and stored at 4C.

To remove any bound nucleic acids, a 5 mL HiTrap Heparin High Performance column (GE Healthcare) was used following a modified protocol (Wong et al., 2009). The protein was first diluted in Heparin buffer (20 mM Tris-HCl, 300 mM NaCl, pH 7.5). The column was preequilibrated with Heparin buffer; the protein was loaded onto the column; the column was washed with 5-10X column volumes of Heparin buffer; and the protein was eluted with a two-step gradient of 700 mM NaCl followed by 1 M NaCl. Fractions containing purified protein were pooled together. To label the protein, a small fraction of protein (~100 ug) was buffer exchanged into PBS and was labelled with the DyLight-594 antibody labelling kit (Thermo Fisher). Glycerol was added to a final concentration of 10% (vol/vol) and aliquots of labelled and unlabeled protein were separately flash frozen in liquid nitrogen and stored at -80C. Specifically, aliquots of 10-20 ul of labelled protein (~1 ug/ul) were frozen while 500 ul of unlabeled protein (~1 ug/ul) were frozen. Prior to buffer exchange, protein was thawed and labelled protein was mixed with unlabeled protein at a ~1:100 mass ratio.

**Bioinformatics**

Disorder predictions on human TFAM were performed using the Predictor of Natural Disordered Regions (PONDR) program (Xue et al., 2010). Images of the crystal structure of TFAM and DNA were obtained from the RCSB Protein Data Bank (rscb.org) (Berman et al., 2006). Proteins involved in mtDNA replication and transcription were selected from the reported mt-nucleoid proteome (Bogenhagen et al., 2008), and sequences were obtained from PubMed based on respective accession numbers. Charged residues were identified using a custom-built Matlab program.

**mtDNA purification**

Total DNA was extracted directly from CRL-1474 cells using a DNeasy Blood & Tissue Kit (Qiagen). To obtain mitochondrial DNA, two overlapping fragments of the mtDNA were amplified by long-range PCR with Takara LA Taq DNA polymerase (Clontech) using two sets of primers (~9 kb fragment: 5′-AACCAAACCCCAAAGACACC-3′ and 5′-GCC AATAATGACGTGAAGTCC-3′ and ~7 kB fragment: 5′-TCCCACTCCTAAACACATCC-3′ and 5′-TTT ATGGGGTGATGTGAGCC-3)(Sundaresan et al., 2015). To directly label mtDNA, trace amounts of fluorescently labelled nucleotides (ChromaTide Alexa Fluor 488 5-dUTP, Thermo Fisher Scientific) were added to the PCR reaction [50 ul reaction containing 10 ng of total DNA, 0.2 uM forward primer, 0.2 uM reverse primer, 1X LA Takara buffer, 400 uM unlabeled dNTPs (each), 2 uM 488-labelled dUTP, 0.5 ul Takara polymerase and PCR-grade water]. Reactions were performed at 94C for 1 min, 30 cycles of 1) 94C at 30s, 2) 55C for 15 s, 3) 68C for 11 min, followed by 72C for 10 min and chilled at 4C. mtDNA fragments were gel extracted after gel electrophoresis and post staining with EtBr. mtDNA gel fragments were further purified with Zymoclean Large Fragment DNA Recovery Kit (Zymo Research) and eluted in low salt buffer (20 mM Tris-HCl, 0 M NaCl, pH 7.5). Unlabeled mtDNA concentration was measured using a Qubit dsDNA HS assay kit (Molecular Probes) on a Qubit Fluorometer, while labelled mtDNA was measured using a DeNovix.

**Nucleic acids**

Single stranded DNA from calf, RNA from calf, RNA from yeast, double stranded DNA from *E. coli*, and double stranded DNA from calf were obtained (Sigma). Free dNTPs (TaKaRa) were mixed with trace amounts of ChromaTide Alexa Fluor 488-5-dUTP (Thermo Fisher Scientific). Nucleic acids and free dNTPs were resuspended in 20 mM Tris, 0 M NaCl, pH 7.5 buffer. To fluorescently label nucleic acids, SYBR Gold (Thermo Fisher Scientific) was added in trace amounts prior to mixing.

**Phase separation assays *in vitro***

Frozen aliquots of protein were thawed at room temperature. Protein solutions were concentrated and buffer exchanged (20 mM Tris-HCl, 500 mM NaCl, pH 7.5) using 0.5 mL Centrifugal filters 3- and 10-kDa (Amicon) using a table top centrifuge at 4C. Final protein concentration was measured by Bradford Assay using BSA standards (BioRad) on a DeNovix spectrophotometer. To induce phase separation, concentrated protein solutions were added to low salt buffer (20 mM Tris-HCl, 0 M NaCl, pH 7.5) to yield final protein concentrations of 1-50 $\mu$M and NaCl concentrations of 150-300 mM. For TFAM-mtDNA experiments, the low-salt solution was prepared first, followed by addition of mtDNA, (final concentration 1-100 ng/$\mu$L), and protein was added last. Solutions were mixed gently and centrifuged prior to imaging.

For high-throughput assays, ~10-50 $\mu$L of protein solution were pipetted to wells of a 384-well plate (PerkinElmer) and covered with 50-100 $\mu$L of mineral oil to prevent evaporation. Pipetting was done either manually or in an automated manner using the ECHO525 liquid handler (Labcyte) into 384 wells. For laser scanning confocal and structured illumination microscopy experiments, 3-6 $\mu$L of solution were added to the center of a 4.5 mm diameter x 0.6 mm depth silicone isolator (Grace-biolabs) on a coverslip and sealed with a glass slide prior to imaging.

Dextran-FITC (3-5 kDa, 40 kDa, 500 kDa, Sigma) solutions were prepared in 1 mg/ml in water. For partitioning experiments, dextran was first was added to the low salt buffer, followed by concentrated protein, to obtain a final concentration of 0.1-0.3 mg/ml. Dextran intensity was measured inside and outside of the droplets to obtain a partition coefficient, which accounted for background intensity. The partition coefficient as a function of dextran size was fit to an exponential to obtain a characteristic length scale.

To capture fusion of droplets, time-lapse movies were performed at two-minute intervals for several fields of view on 384-well plates using high-throughput confocal microscopy immediately after mixing. Fusion events captured within the first 1-2 hours were manually identified, and the aspect ratio as a function of time was quantitatively analyzed using Matlab. The characteristic time scale, $\tau$, and the size of the drop, $\ell$, were used to estimate the inverse capillary velocity, $\tau/\ell = \eta/\Upsilon$ (Brangwynne et al., 2011).

To determine the mobility within the droplets, a ~1 um spot was photobleached precisely 30 minutes after mixing. The fraction recovered was measured as a function of time and normalized to obtain a characteristic time scale, $\tau$ (Phair et al., 2003).

**Live-cell imaging**

For live-cell imaging, primary skin fibroblasts and HeLa cells were seeded 1-3 days prior to the experiment in 8-well imaging chambers (Thermo Fisher Scientific). Cells were washed a few times with pre-warmed complete media lacking phenol red to replace the original cell media prior to imaging. For FRAP experiments, HeLa cells expressing TFAM-mKate2 were incubated with PicoGreen 0.3% (vol/vol) (Ashley et al., 2005) and 100 nM MitoTracker Deep Red at least one hour prior to imaging on the LSM780. For coarsening experiments, HeLa and primary HGPS cells were also pretreated with MitoTracker Deep Red (100 nM), PicoGreen (0.3% vol/vol) for ~ 1 hour, and/or with Ethidium Bromide (1 ug/ml) for ~0.5-1 hour prior to imaging with either laser scanning confocal microscopy or Airyscan confocal microscopy (Ashley and Poulton, 2009). To osmotically induce mitochondrial swelling, normal media was replaced with DI water and live cells were immediately imaged.

**Immunofluorescence**

Cells were seeded in 96-well or 384-well plates (PerkinElmer) or in 12-well dishes 24 hours prior to fixation. To label the mitochondrial network, cells were incubated with 100 nM MitoTracker Red/Deep

Red (Thermo Fisher Scientific) for 15-30 min at 37C. Cells were immediately washed with PBS and fixed with 4% PFA for 10 min. Cells were washed and permeabilized with 0.1% Triton-X for 10 min. Cells were washed and incubated with primary antibodies in PBST with 5% BSA for 1 hour at room temperature or overnight at 4C. The following antibodies were used: anti-DNA (EMD Millipore, clone AC-30-10), anti-TOMM20 (Sigma, HPA011562), anti-TFAM (Sigma, HPA063684), anti-TFB2M (Sigma, HPA028482), anti-SSBP1 (Sigma, HPA002866), anti-TOP1MT (Sigma, HPA001915), anti-HSPD1 (Sigma, HPA001523), anti-LONP1 (Proteintech, 154401-A), anti-CLPP (Thermo Fisher Scientific, PA5-52722), anti-HSP10/EPF (RND Systems, MAB3298), anti-mtHSP70 (Thermo Fisher Scientific, MA3-028). Cells were washed 1X with PBST and 2X with PBS followed by incubation with secondary antibodies for 1 hour at room temperature. The following secondary antibodies were used: anti-rabbit (Thermo Fisher Scientific, as purchased) and anti-mouse (Thermo Fisher Scientific, as purchased). Cells were washed 1X with PBST and 2X with PBS followed by an incubation with DAPI for 30 minutes. Cells were stored in PBS at 4C. 96-well plates were sealed for use in high-throughput imaging, while coverslips were mounted onto glass slides with VectaShield mounting medium (Vectorlabs) and sealed with nail polish or with ProLong Gold Antifade Mountant (Thermo Fisher Scientific), as purchased, and left to cure for >24 hours for use in super-resolution imaging. For all high-throughput microscopy, all immunofluorescent experiments had three technical replicates with >4 fields of view per well and three experimental replicates. For super-resolution imaging, at least two experimental replicates were performed on three or more cells per condition.

**RNA FISH**

Custom Stellaris® FISH Probes were designed against mt-12S and mt-COI by utilizing the Stellaris® RNA FISH Probe Designer (Biosearch Technologies, Inc., Petaluma, CA) available online at www.biosearchtech.com/stellaris designer (version 4.2). Primary cells in 96-well plates or in 12-well plates were hybridized with the mt-12S and mt-COI Stellaris RNA FISH Probe set labeled with Qusar-670 (Biosearch Technologies, Inc.), following manufacturer's instructions available online at www.biosearchtech.com/stellarisprotocols for sequential immunofluorescence and RNA FISH. Briefly, cells were fixed in 4% PFA for 10 min and permeabilized with 0.5% Triton-X for 10 min. Immunofluorescence was performed first without blocking agents and cells were fixed again for 10 min with 4% PFA. Manufacturer's instructions were followed for subsequent FISH labelling and samples were stored in PBS.

**BrU incorporation**

Cells were incubated with 2.5 mM BrU (Sigma) for 1 hour in cell media. Media was removed and replaced with media containing 100 nM MitoTracker Red for 15-30 minutes to label the mitochondrial network and remove any diffuse BrU (Jourdain et al., 2013; Ramos et al., 2019). Cells were subsequently washed briefly with PBS, fixed 10 min with PFA and permeabilized with 70% ethanol for 1 hour at 4C followed by 10 min of 0.5% Triton X at room temperature. Primary antibodies for anti-BrdU (Sigma, 11170376001) and anti-TFAM (Sigma, HPA063684) were applied for 1 hour at room temperature followed by secondary antibodies without the use of blocking agents.

**Light microscopy techniques**

High-throughput confocal imaging was performed on fixed cells in 96- and 384-well plates (PerkinElmer) and on phase separation assays on 384-well plates (PerkinElmer) using the fully-automated Yokogawa CV7000S spinning disk microscope using a 60X water immersion objective with 405, 488, 561 and 647 nm laser lines as well as with brightfield imaging. Images were acquired as maximum intensity projections of 6-10 z-slices with 1 um thickness.

Structured Illumination Microscopy (SIM) was performed on fixed cells and TFAM-mtDNA droplets using ELYRA PS.1 on an AxioObserver Z1 inverted microscope controlled by ZEN software. A 63X/1.4 NA oil Plan Apochromat objective was used with 405, 488, 561, and 647 nm laser lines.

Laser scanning confocal microscopy was performed using a Carl Zeiss LSM780 inverted microscope controlled by ZEN software and with definite focus. Live cells were imaged using either 63X/1.46 NA oil objective and a 100X/1.46 NA objective for FRAP experiments with 488, 561, and 633 nm laser lines using a $CO_2$/heating control stage insert. TFAM phase separated droplets were imaged on a 60X objective with 594 laser lines and in brightfield/DIC. For *in vitro* FRAP experiments, a ~1 um spot corresponding to TFAM-Dylight 594 and/or mtDNA-Alexa488 was bleached using 488 and 561 nm light within the center of the droplet using maximum laser power and recovery was monitored over a period of 10-30 minutes. For *in vivo* FRAP experiments, a ~0.5 um spot corresponding to TFAM-mKate2 was bleached using 488 and 561 nm light inside mt-nucleoids using maximum laser power and recovery was monitored over a period of ~10 minutes.

Airyscan confocal microscopy was performed using a Carl Zeiss LSM880 inverted confocal microscope controlled by ZEN software and with definite focus. Coarsening experiments in live cells were imaged using a 63X/1.46 NA oil objective with 488, 561, and 633 laser lines and inside of an incubation chamber.

**Image analysis**

Images and time-lapse movies were visualized and created using Fiji, Zen, Matlab, and Imaris software. For representation in figures, some images were filtered using standard Gaussian, mean and/or band pass filters. For Figures 1D-1F, 1H-1I, 1J-1K, 2B, 2C, 2D, 3C-3D, 3H-3I, 4A-4B, 4C, 4E, and 6A-6E the same brightness/contrast settings were used across conditions from the same experiment. For Figures 1A-1B, 2G, 3E-3G, and 5D-5E, the same brightness/contrast settings were not used, but rather adjusted for each panel for clarity.

All quantitative image analysis was performed using custom-built Matlab code. Image analysis of phase separated droplets consisted of measuring the average intensity, size, aspect ratio, and number of droplets per field of view. These values were averaged for ≥4 fields of view per well.

For analysis of high-throughput images, nuclear markers, such as DAPI, were used to identify individual cells while cytoplasmic markers, such as MitoTracker Red, were used to segment cells using a watershed algorithm. To quantify the extent of morphological damage to the nucleus, the mean negative curvature of the nucleus was obtained for each cell following a published method (Driscoll et al., 2012), and mean intensity within the nucleus was also computed. To detect damaged mitochondria, the mitochondrial network was filtered with a Gaussian filter, segmented and thinned. The mitochondria within each cell were analyzed based on their size, brightness, and shape to identify the phenotype of interest: swollen, bright mitochondria. The number of damaged mitochondria per cell was averaged across all cells within a well. All wells were averaged per technical replicate (total of three technical replicates per condition), and then averaged for all experimental biological replicates, where n=3.

Within each cell, individual mitochondrial nucleoids labeled with anti-DNA were detected and analyzed using adapted Matlab Multiple Particle Tracking Code adapted (see http://physics.georgetown.edu/matlab/index.html) (Crocker and Grier, 1996). The coordinates of identified mt-nucleoids were mapped onto the channel containing signal of nucleoid protein markers, such as TFAM, to correlate integrity intensity of anti-DNA with that of the associated protein. Similar analysis was performed to quantify RNA transcript levels, where nucleoids were identified with anti-TFAM that were later correlated with the intensity in the channel labelled by FISH or anti-BrU. The nucleoids were then mapped on to the mitochondrial network to identify if they were in a normal or damaged mitochondrion. Similar analysis was performed on $UPR^{mt}$ markers, except mean intensity of the $UPR^{mt}$ marker for each segmented mitochondrion was used instead of intensity at the nucleoid. On a per cell basis, the average intensity of nucleoid markers from damaged mitochondria was compared to the average intensity of nucleoid markers from undamaged mitochondria. Average values were computed for each technical replicate and then averaged for all experimental replicates. Intensity values were normalized to the average intensity values of the WT-1 cell line.

Colocalization of mutants was quantified using Pearson's correlation coefficient for specified pairs of channels using intensity values for all pixels within the brightest z-plane for each segmented nucleoid (Dunn et al., 2011). Correlation coefficients were averaged for all nucleoids (n≈20-40) in 4-5 cells per construct.

**Western Blot**

Cells were collected, washed in PBS, dissolved in 2X Laemmli Sample Buffer (Bio-Rad) and denatured for 5-10 min at 95C and stored at -20C. Samples were loaded onto a 10-well 4-12% Bis-Tris Protein Gel at 150V for 1 hour and transferred to a membrane via wet transfer at 250 mA for 90 min. Antibodies used were Lamin A/C (Santa Cruz, 376248), β-actin (Sigma, A2228), TFAM (Sigma, HPA063684), HSPD1 (Sigma, HPA001523), and ATF5 (Abcam, ab60126). Western blots were performed on three-six independent experimental replicates and band intensities were quantified using Bio-Rad Image Lab software.

**mtDNA copy number**

Total DNA (both genomic and mitochondrial) was purified using the DNeasy Blood and Tissue Kit (Qiagen). DNA concentration was measured using a Denovix. qPCR was performed on 50 ng of DNA in a 20 ul mixture on 96 well plates using a CFX Real-Time PCR instrument (Bio-Rad). Primers for mtDNA $tRNA^{Leu(UUR)}$ (5'- CACCCAAGAACAGGGTTTGT-3' and 5' TGGCCATGGGTATGTTGTTAA-3') and nuclear 18S rDNA (5' TAGAGGGACAAGTGGCGTTC-3', 5'-CGCTGAGCCAGTCAGTGT-3') were used as previously described (Xiong et al., 2016). Six independent experimental replicates were performed, each with three technical replicates. Bio-Rad CFX maestro software was used to determine the average $C_t$ for each experiment. The relative mtDNA copy number was obtained as $2 \times 2^{\Delta C_t}$ where $\Delta C_t = C_t^{nuclear\ DNA} - C_t^{mtDNA}$ (Gonzalez-Hunt et al., 2016).

**Estimating concentration of TFAM and DNA in a single mitochondrion**

A single mitochondrial nucleoid has a diameter of ~100 nm and is estimated to have 1 molecule of mtDNA and ~1,000 molecules of TFAM (Kukat et al., 2011). Based on these measurements, the volume of a spherical mitochondrial nucleoid is ~$5\times10^{-22}$ $m^3$ or ~$5\times10^{-19}$ L, which yields a concentration of approximately $C_{TFAM}^{nucleoid}$ ≈3 mM or ~100 μg/μl. Mitochondrial DNA is roughly 16 kb, which has a molar mass of ~10 Mg/mol, which yields $C_{mtDNA}^{nucleoid}$ ≈ 3 μM or 30 μg/μl inside a mitochondrial nucleoid. This gives a molar ratio of ~0.001 and a mass ratio of DNA/TFAM ≈ 0.3 within the nucleoid. To determine the diffuse concentration of mtDNA and TFAM if a single mitochondrial nucleoid was fully solubilized in the mitochondrion, we assume a unit mitochondrion to be a cylinder with 1 $\mu$m in length and 0.2 $\mu$m in radius, which gives a unit volume of ~1x$10^{-19}$ $m^3$ or ~1x$10^{-16}$ L. Assuming all 1,000 molecules of TFAM and 1 molecule of mtDNA are uniformly distributed in the unit mitochondrion, we obtain concentrations of $C_{TFAM}^{mitochondria}$ ≈ 20 µM or 0.5 µg/µl and $C_{mtDNA}^{mitochondria}$ ≈15 nM or 0.15 µg/µl, which also gives a molar ratio of ~0.001 and a mass ratio of DNA/TFAM ≈ 0.3 that is comparable to the conditions used in our in vitro experiments.

**qPCR analysis**

RNA was isolated from primary cells using a RNeasy Plus Micro Kit (Qiagen), and cDNA was obtained from 1 ug of RNA after reverse transcription using an iScript cDNA Synthesis Kit (Bio-Rad). 10 ul RT-PCR reactions containing a SYBR Green Master Mix (Bio-Rad) were dispensed using an Echo 525 Liquid Handler (Labcyte) into Hard-Shell 384-well PCR plates (Bio-Rad). qPCR was performed on a CFX384 Touch Real-Time PCR Detection System (Bio-Rad) and Ct values were obtained using CFX Maestro Software (Bio-Rad). Each qPCR reaction had three technical replicates, and data were averaged for three biological replicates (Taylor et al., 2019). 12S and mtCOI transcripts were analyzed relative to ACTB, ATF5 was normalized relative to TUBB1, and all other $UPR^{mt}$ transcripts were analyzed relative to TBP. Primers are listed in Extended Data Table 1.

**Mitochondrial oxygen consumption**

Cells were plated at 50,000/well in 96-well Seahorse cell culture plate containing DMEM media with 15% FBS, 2 mM GlutaMAX and 1 mM Sodium Pyruvate (Gibco). Cells were incubated overnight for 18 hours before media was replaced with Seahorse base XF medium containing 2 mM GlutaMAX, 1 mM Sodium Pyruvate, and 25mM Glucose, at pH 7.4. Cell plates were then placed into a non-CO2 incubator at 37°C for 1 hour for equilibration before running the XF96 Extracellular Flux Analyzer standard mitochondrial stress test. The three injection drugs were administered as follows: Port A) Oligomycin: 2.5 uM; Port B) FCCP 2 uM; Port C) Antimycin A 2 uM. In our studies, the outermost wells in each 96-well plate were not used as a precaution against hypothetical temperature inequality/edge effects. At the end of seahorse run, the medium was removed, and 50 ul of 1x RIPA lysis buffer was added to each well, the plate was frozen at -20C, then thawed at 4C on a shaker for 1 hour, the lysate was collected into 1.5 ml microcentrifuge tube, spun down, and the supernatant was collected to measure protein concentration by standard BCA method. The determined protein concentrations from each well were used to normalize oxygen consumption rates.

**Mitochondrial membrane potential and reactive oxygen species generation**

Mitochondrial membrane potential (MMP) and mitochondrial reactive oxygen species (ROS) were measured by a BD FACSCanto II flow cytometer. HGPS cells were plated in 6-well plate at a confluence of 200K cells/well. On the day of experiment, cells were washed in PBS and harvested by trypsinization, washed once more with PBS, and resuspended in DMEM without phenol red (Invitrogen). Cells were incubated with dyes Tetramethylrhodamine, methyl ester (TMRM) (40 nM for 15 min) to detect MMP, and mitoSOX (3 μM for 30 min) for mitochondrial ROS (all from Life Technologies), fluorescence was determined with flow cytometry from $1X10^4$ cells. Data were analyzed using FCS Express 4 software.

**Statistical methods**

For high-throughput imaging experiments with fixed cells, three technical replicates were performed, each with ≥5 fields of view, and averaged to obtain the measurement for that experiment. These experiments were repeated independently three times (n=3) and the values for each experiment were averaged. For high-throughput experiments with in vitro droplets, each condition was assayed per well, with ≥9 fields of view. Measurements from each field of view were averaged together per experiment, and each experiment was repeated independently at least three times. For all experiments, measurements were repeated on ≥3 days. For FRAP experiments, all FRAP measurements were pooled together and averaged. For mitochondrial functional testing, three experimental replicates (each with technical replicates) were performed for each cell line and averaged. The data for WT and HGPS cell lines were pooled for some measurements as indicated. For all experiments, error was reported as standard deviation or standard error of the mean as indicated. Number of damaged mitochondria, mitochondrial membrane potential, and mitochondrial ROS were compared using a single-factor analysis of variance (ANOVA) statistical test and the p-value for the test statistic was determined using Excel and R and was reported in the figure legend. Pairs of means were determined statistically significant using the two-sided Fisher Least Significant Difference (LSD) method and indicated graphically. P-values were determined for Basal Respiration, Maximal Respiratory Capacity and Reserve Capacity using a two-sided Student's t-test based on the standard deviation from experimental replicates.

**Code availability**: All code used in this study can be available from the corresponding author on request.

**Supplementary Table S1: qPCR primers**

| qPCR target | Forward Primer | Reverse Primer |
|---|---|---|
| 12S* | 5’-ATGCAGCTCAAAACGCTTAGC-3’ | 5’-GCTGGCACGAAATTGACCAA-3’ |
| mtCOI* | 5’-CAGCAGTCCTACTTCTCCTATCTCT-3’ | 5’-GGGTCGAAGAAGGTGGTGTT-3’ |
| ACTB | 5’-CATGTACGTTGCTATCCAGGC-3’ | 5’-CTCCTTAATGTCACGCACGAT-3’ |
| ATF5† | 5’-CTGGCTCCCTATGAGGTCCTTG-3’ | 5’-GAGCTGTGAAATCAACTCGCTCAG-3’ |
| TUBB1 | 5’-CTACAACGCGGTTCTGTCTATC-3’ | 5’-GGTGGGTGTCGTCAGCTTC-3’ |
| CLPP‡ | 5’-GCGCGCCTATGACATCTACT-3’ | 5’-AACGCTGTCATCGATCGGG-3’ |
| LONP1‡ | 5’-CATGACGATCCCCGATGTGT-3’ | 5’-ACATCCGACTCATTGCTGTCA-3’ |
| HSPE1 (mtHSP10)‡ | 5’-AGTAATGGCAGGACAAGCGT-3’ | 5’-ACTGGTTGAATCTCTCCACCC-3’ |
| HSPD1 (mtHSP60)‡ | 5’-TGCTTCGGTTACCCACAGTC-3’ | 5’-ACTGTTCTTCCCTTTGGCCC-3’ |
| HSPA9 (mtHSP70)‡ | 5’-ACAAGCAAAGGTGCTGGAGA-3’ | 5’-GGCATTCCAACAAGTCGCTC-3’ |
| TBP | 5’-GAGCCAAGAGTGAAGAACAGTC-3’ | 5’-GCTCCCCACCATATTCTGAATCT-3’ |

*Primers normalized to ACTB. †Primers normalized to TUBB1. ‡Primers normalized to TBP.

### Supplementary Figure S1

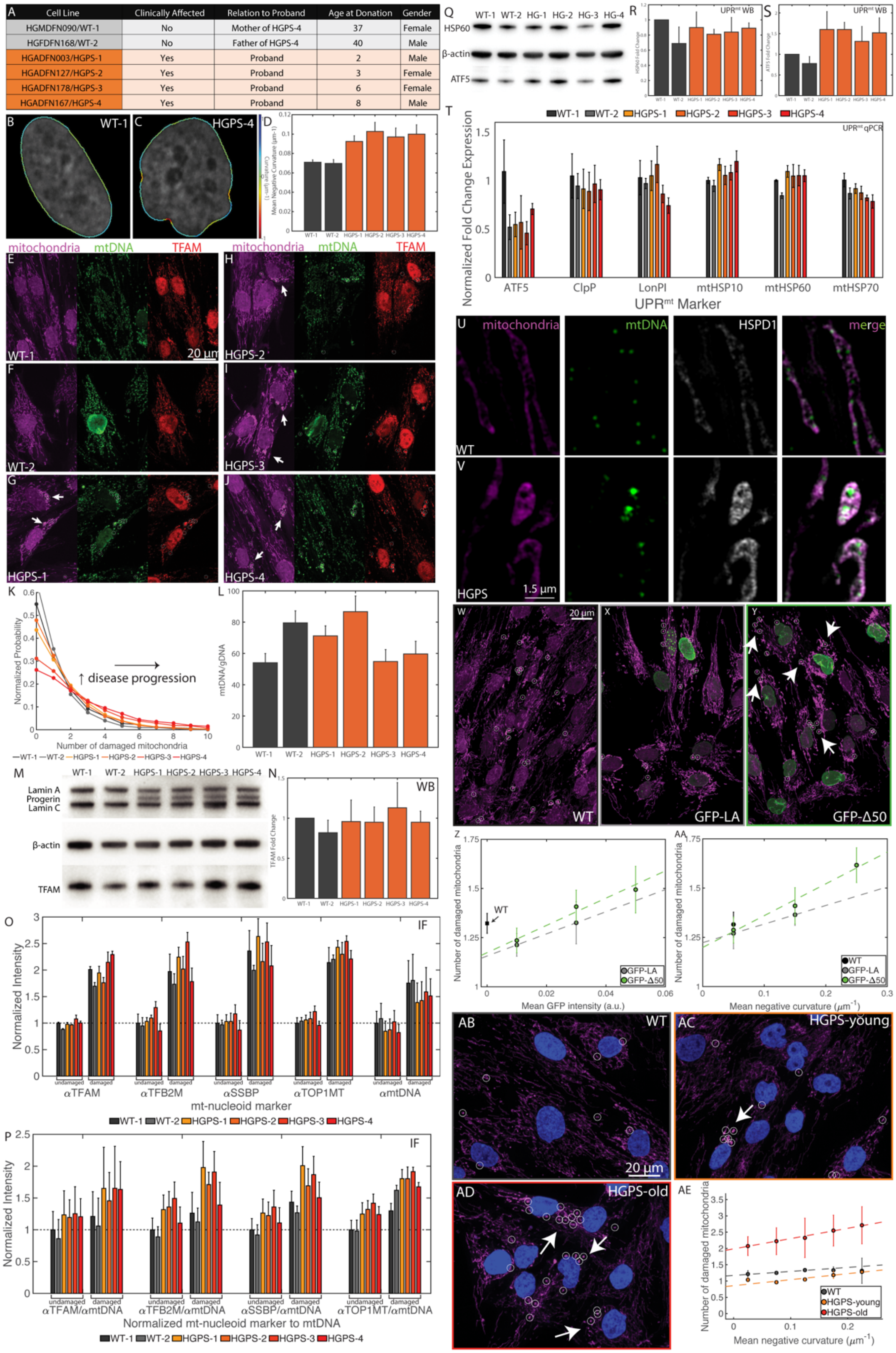

**Fig. S1:** (A) Table summarizing primary skin fibroblast cell lines obtained from the Progeria Research Foundation (PRF). Categories included if patients are clinically affected with HGPS, their relation (if any), age at donation, and gender. (B-C) Images of typical segmentation of a nucleus exhibiting the normal, smooth phenotype from WT-1 cell line (B). or the irregular morphology from HGPS-4 cell line (C). Color scale shows curvature along the nuclear perimeter, where red = -1 and blue = 1. (D) Average mean negative curvature for all cell lines. Error bars represent standard deviation (where n =3 experimental replicates, each with 8 technical replicates containing 5 fields of view. Approximately ~700-3,000 cells analyzed per experimental replicate). (E-J) Representative high-throughput confocal images of WT-1 (E), WT-2 (F), HGPS-1 (G), HGPS-2 (H), HGPS-3 (I), and HGPS-4 (J) cell lines. Left image is mitochondria (MitoTracker Red, magenta); center image is mtDNA (anti-DNA, green); right image is TFAM (anti-TFAM, red). Scale bar = 20 μm. (K) Normalized probability distribution of number of damaged mitochondrial nucleoids for all cells of each primary cell line, which corresponds to data shown in Fig. 1C. (L) qPCR results of mtDNA copy number relative to genomic DNA for all cell lines tested, n = 6 experimental replicates and data are reported as mean±standard error. (M) Western blot results for all six primary cell lines showing bands for Lamin A, Progerin, and Lamin C as well as TFAM with B-actin as the housekeeping protein. (N) Quantification of TFAM protein levels from Western Blot in M, where n = 6 experimental replicates and data are reported as mean±standard error. (O) Based on high-throughput imaging, integrated intensity values for markers for nucleoid associated proteins (anti-TFAM, anti-TFB2M, anti-SSBP, and anti-TOP1MT) and anti-DNA were compared for nucleoids from undamaged and damaged mitochondria analyzed for each cell and averaged for all cells, where n=3 experimental replicates and error bars represent standard error. (P) Average ratio of normalized nucleoid intensities from undamaged and damaged mitochondria on a per cell basis as a function of the mitochondrial marker (anti-TFAM, anti-TFB2M, anti-SSBP, and anti-TOP1MT). Normalization was done by taking the ratio of the integrated intensities of the protein marker over the respective mtDNA signal for each nucleoid, where n=3 experimental replicates and error bars represent standard error.

(Q) Western blot results for all six primary cell lines showing bands for $UPR^{mt}$ markers anti-HSPD1 (mtHSP60) and ATF5 with B-actin as the housekeeping protein. (R) Quantification of HSPD1 (mtHSP60) protein levels from Western Blot in Q, where n = 3 experimental replicates and data are reported as mean±standard error. (S) Quantification of ATF5 protein levels from Western Blot in Q, where n = 3 experimental replicates and data are reported as mean±standard error (p=0.3). (T) qPCR results of $UPR^{mt}$ markers (ATF5, ClpP, LonPI, mtHSP10, mtHSP60, mtHSP70) in all six primary cell lines reported as normalized fold change expression, where n=3 independent experimental replicates, each with three technical replicates, and error bars are standard error. (U-V) Representative z-slices of SIM images of WT (U) and HGPS (V) cell lines. Left image is mitochondria (MitoTracker Red, magenta); mtDNA (anti-DNA, green); HSPD1/mtHSP60 (anti-HSPD1, gray); right image is the merged image. Scale bar = 1.5 µm. (W-Y) High-throughput confocal images of mitochondria from WT immortalized cells (W), GFP-lamin A (X), and GFP-progerin (Y) with labelled mitochondria (MitoTracker Red, magenta) and GFP (green). Arrow heads point to GFP-progerin cells that exhibit the damaged mitochondrial phenotype observed in advanced HGPS cells. White circles indicate damaged mitochondria detected computationally. Scale bar = 20 µm. (Z-AA) Number of damaged mitochondria as a function of mean GFP intensity (Z) and mean negative curvature (AA) for WT (black), GFP-lamin A (gray), and GFP-progerin (green). Cells were binned into equal bins, containing >50 cells per bin. Error bars are standard error for n = 3 experimental replicates, each with 9 technical replicates containing >16 fields of view (~600-3000 cells per experimental replicate for each cell line). Linear weighted fits are shown by dashed line. (AB-AD) High-throughput confocal images of mitochondria from WT primary cells (AB), HGPS-young primary cells (AC), and HGPS-old primary cells (AD) with labelled mitochondria (MitoTracker Red, magenta) and nuclei (DAPI, blue). Arrow heads point to HGPS cells that exhibit damaged mitochondrial phenotype. White circles indicate damaged mitochondria detected computationally. Scale bar = 20 µm. (AE) Number of damaged mitochondria as a function of mean negative curvature primary cells analyzed in Fig. 1. Cells were binned into equal bins, containing >50 cells per bin. Error bars are standard error for n = 2 experimental replicates, where primary cell lines have been grouped: WT (WT-1 and WT-2), HGPS-young (HGPS-1 and HGPS-2) and HGPS-old (HGPS-3 and HGPS-4). Each cell line has 8 technical replicates containing >5 fields of view (1500-4000 cells per experimental replicate for each cell line). Linear weighted fits are shown by dashed line.

**Supplementary Figure S2:**

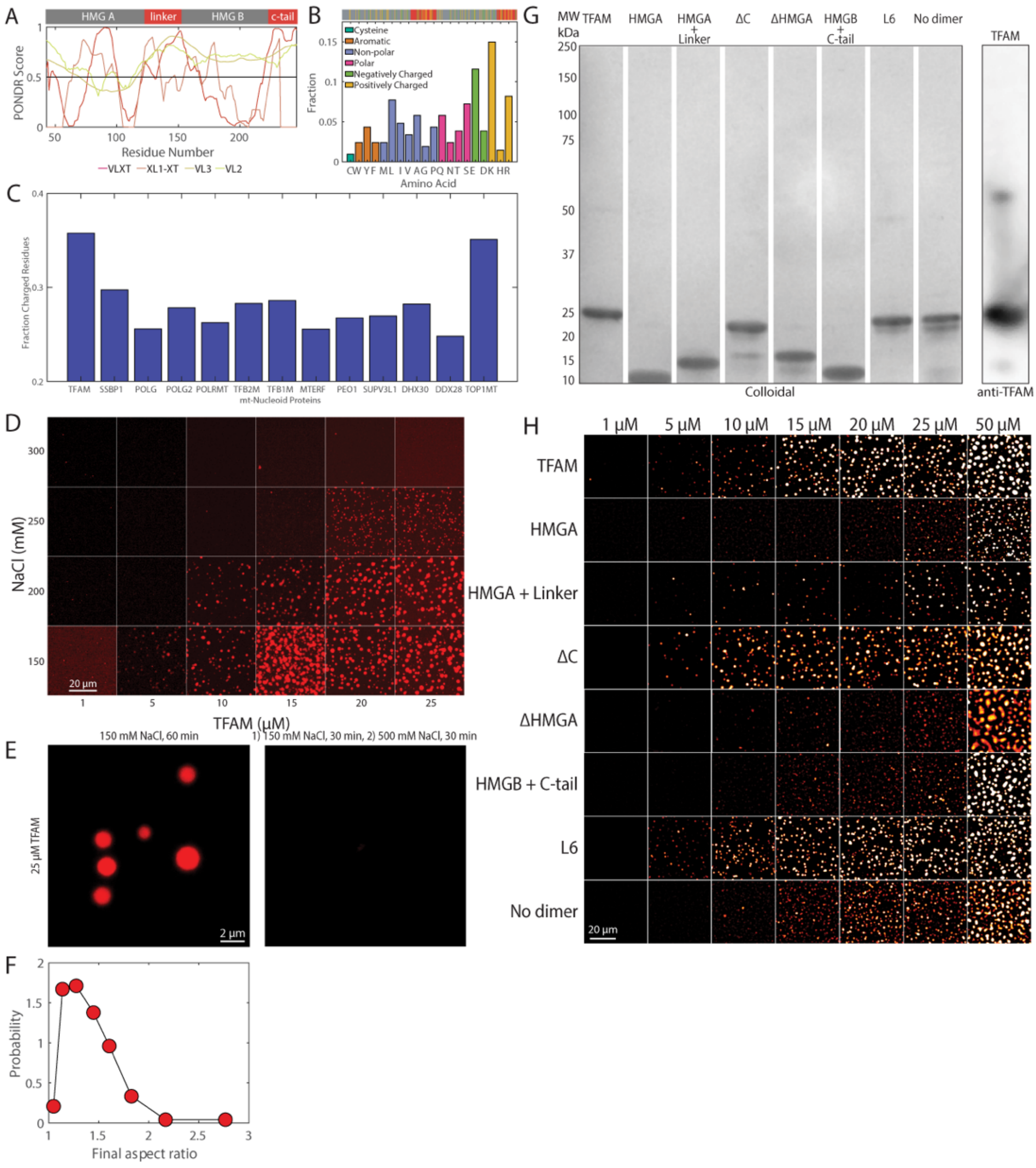

**Fig. S2:** (A) PONDR score as a function of residue position of full length TFAM using the VLXT, XL1-XT, VL3, and VL2 algorithms. Top protein domain diagram shows corresponding functional domains with HMGs in gray and disordered domains in red. (B) Frequency of amino acids in full length TFAM. Top protein domain diagram shows position of negatively charged (green) and positively charged (yellow) residues. (C) Fraction of charged residues of core proteins in the mitochondrial nucleoid proteome involved in replication and transcription. (D) Images from a representative high-throughput microscopy experiment used to generate the phase diagram in Fig. 2A. Scale bar = 20 μm. (E) Reversibility of phase separation experiment. Left panel shows contents after 60 minutes post mixing at 25 μM TFAM and 150 mM NaCl (Sample 1). Right panel shows contents from Sample 1 30 minutes after mixing followed by an additional 30 minutes in high salt buffer to reach 25 μM TFAM in 500 mM NaCl (Sample 2). Scale bar = 2 μm. (F) Probability distribution of the aspect ratio upon completion of the fusion events observed in Fig. 2F. (G) Protein purification results. Left panel shows gel using Colloidal Blue Stain of purified full length TFAM and mutants. Right panel shows Western blot using anti-TFAM to confirm specificity of protein. (H) Images from high-throughput microscopy experiment used to generate the phase diagram and images in Fig. 2F,2G for TFAM and mutants ranging in concentration from 1, 5, 10, 15, 20, 25 and 50 μM. Scale bar = 20 μm.

**Supplementary Figure S3:**

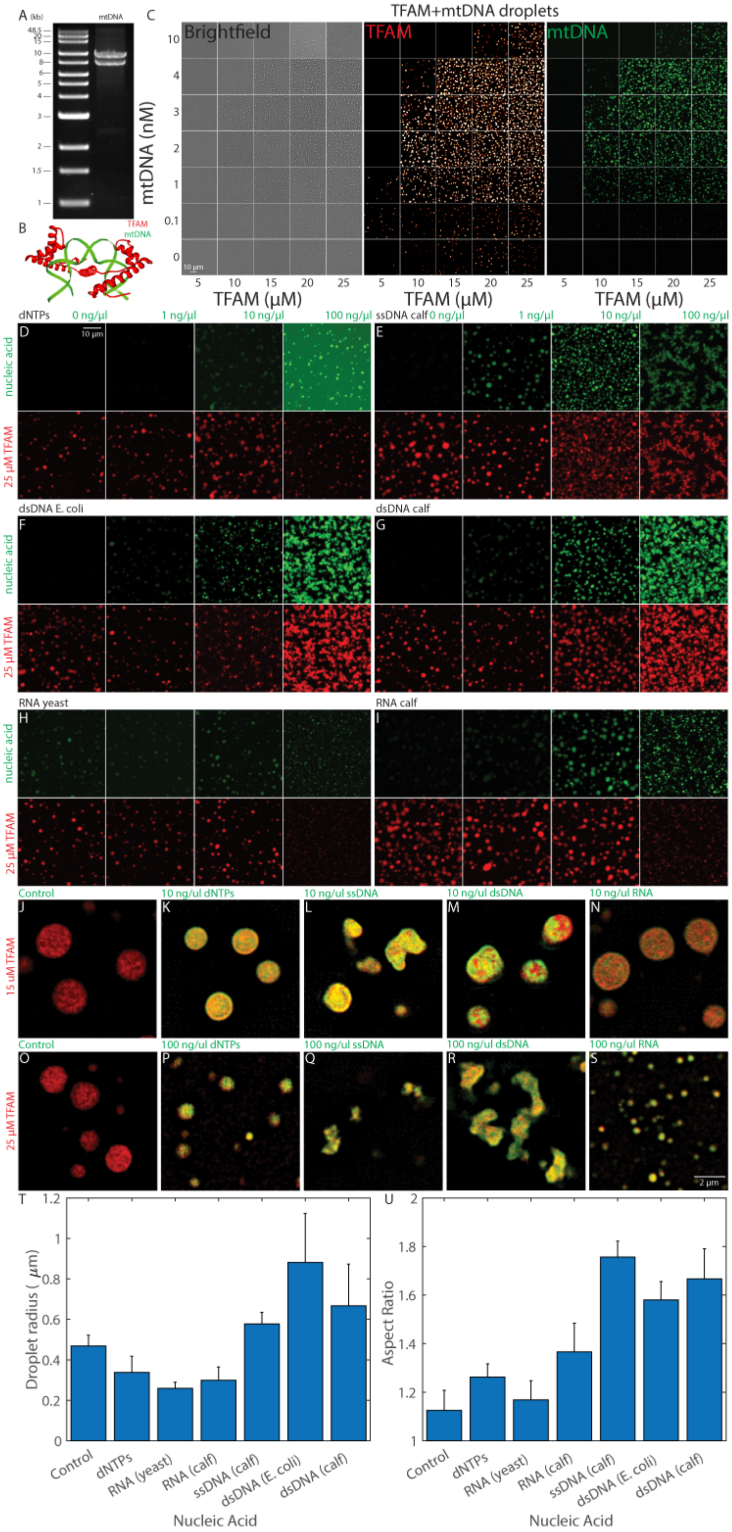

**Fig. S3:** (A) Agarose gel stained with Ethidium Bromide showing purification of two mtDNA fragments (~7 and ~9 kb) after long-range PCR. (B) Cartoon illustrating TFAM-mtDNA binding based on structure obtained from X-ray crystallography. Adapted from Protein Data Base (see Methods). (C) An example high-throughput imaging assay of droplet formation upon various TFAM and mtDNA concentrations, indicated by brightfield, TFAM localization in red heat map, and mtDNA localization in green (labeled with dUTP-Alexa488). (D-I) High-throughput images of nucleic acid addition to TFAM droplets. 25 μM TFAM (final) was mixed with nucleic acids containing 0, 1, 10, and 100 ng/μl (final) of dNTPs (D), ssDNA from calf (E), dsDNA from E. Coli (F), dsDNA from calf (G), RNA from yeast (H), and RNA from calf (I). Top row is of the nucleic acids labelled with SYBR Gold or of dNTPs with dUTP-Alexa488, and bottom row contains droplets labelled with TFAM-Dylight594. Scale bar = 10 μm. Intensities were matched across the conditions within each panel. (J-S), SIM images of TFAM (red) droplets mixed with various nucleic acids (green). (J-N) are at 15 μM TFAM, where J = control, K = 10 ng/μl dNTPs, L= 10 ng/μl ssDNA calf, M = 10 ng/μl dsDNA calf, and N = 10 ng/μl RNA calf. (O-S) 25 μM TFAM, where O = control, P = 100 ng/μl dNTPs, Q = 100 ng/μl ssDNA calf, R = 100 ng/μl dsDNA calf, and S = 100 ng/μl RNA calf. Scale bar = 2 μm. (T) Droplet radius as a function of nucleic acid added, where n = 3 experimental replicates and error bars represent standard error of the mean. (U) Aspect ratio of droplets as a function of nucleic acid added, where n = 3 experimental replicates and error bars represent standard error of the mean.

**Supplementary Figure S4:**

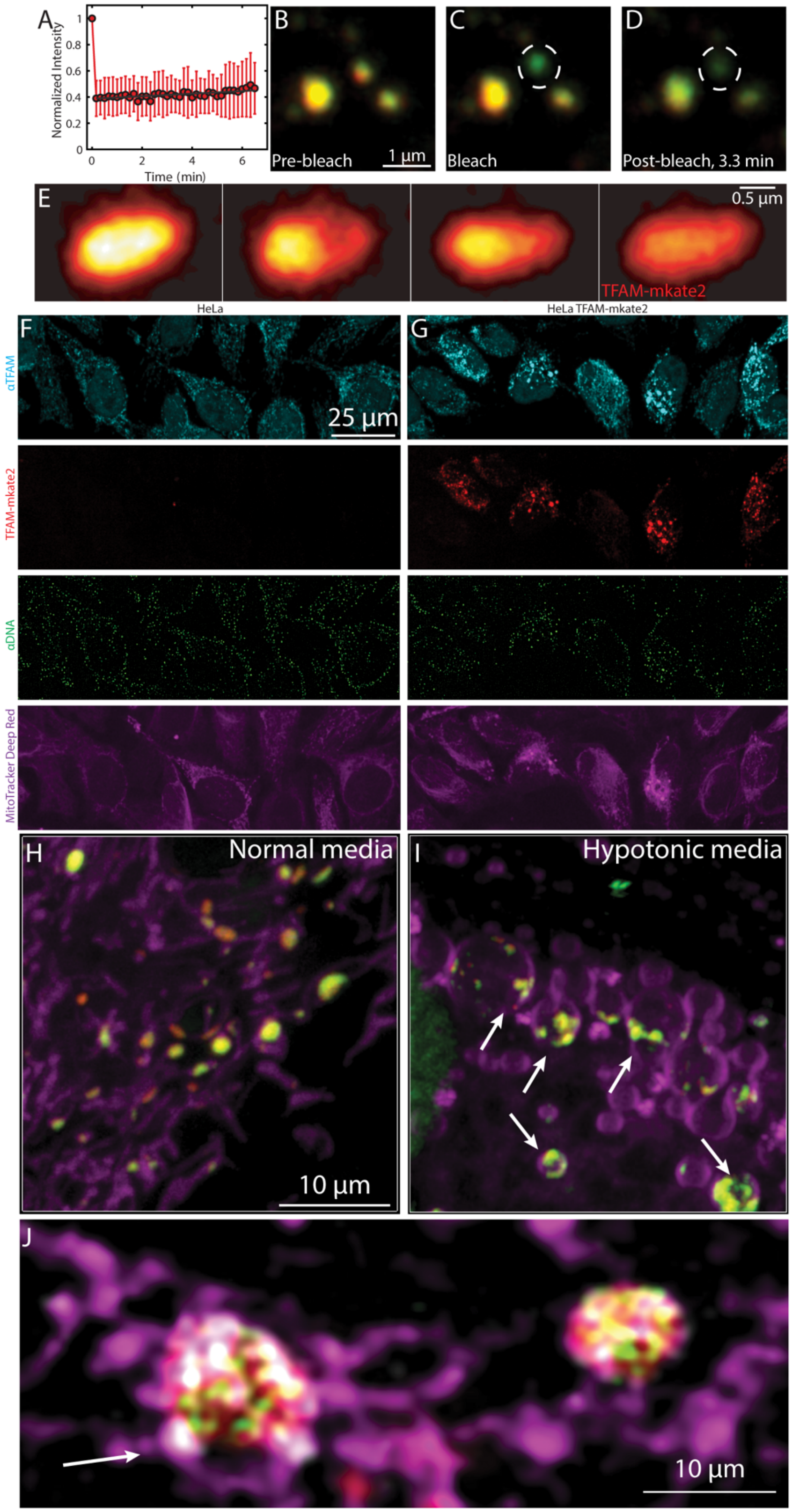

**Fig. S4:** (A) FRAP recovery as a function of time on TFAM-mKate2 in live HeLa cells (n=20 cells, error bars represent standard deviation). For these experiments, bleaching (dashed circle) was performed on small nucleoids as to bleach the entire structure. (B-D) Images from FRAP experiment, showing pre-bleach (B), bleach (C), and post-bleach after 3.3 minutes (D). Images are an overlay of TFAM-mKate2 (red) and PicoGreen (green). Dashed circle indicates which nucleoid was bleached. Scale bar = 1 µm. (E) The same images as in Fig. 4C, except the intensity of only the TFAM-mKate2 channel is shown using a hot heatmap. (F-G) High-throughput confocal images of Hela cells (F) and Hela cells over-expressing a TFAM-mKate2 construct (G), where (n= 1 experimental replicate, with 3 technical replicates each with 16 fields of view). Cells were labelled as anti-TFAM (cyan), TFAM-mKate2 (red), anti-DNA (green), and MitoTracker Deep Red (magenta). Scale bar = 25 µm. (H-I) Images of mitochondria (MitoTracker Deep Red, magenta), TFAM (TFAM-mKate2, red), and mtDNA (PicoGreen, green) in HeLa cells in normal media (H) and hypotonic media (water) (I). Arrow heads point to nucleoids that remain as discrete structures in swollen mitochondria. Scale bar = 10 µm. (J) Single z-slice of a SIM image of enlarged nucleoids in fixed HeLa cells, where TFAM (TFAM-mKate2, red), mtDNA (anti-DNA, green), (anti-TFB2M, gray), and mitochondria (MitoTracker Deep Red, magenta) are labelled. Arrow points to swollen mitochondria with nucleoids occupying a fraction of the mitochondrial matrix. Scale bar = 10 µm.

**Supplementary Figure 5**

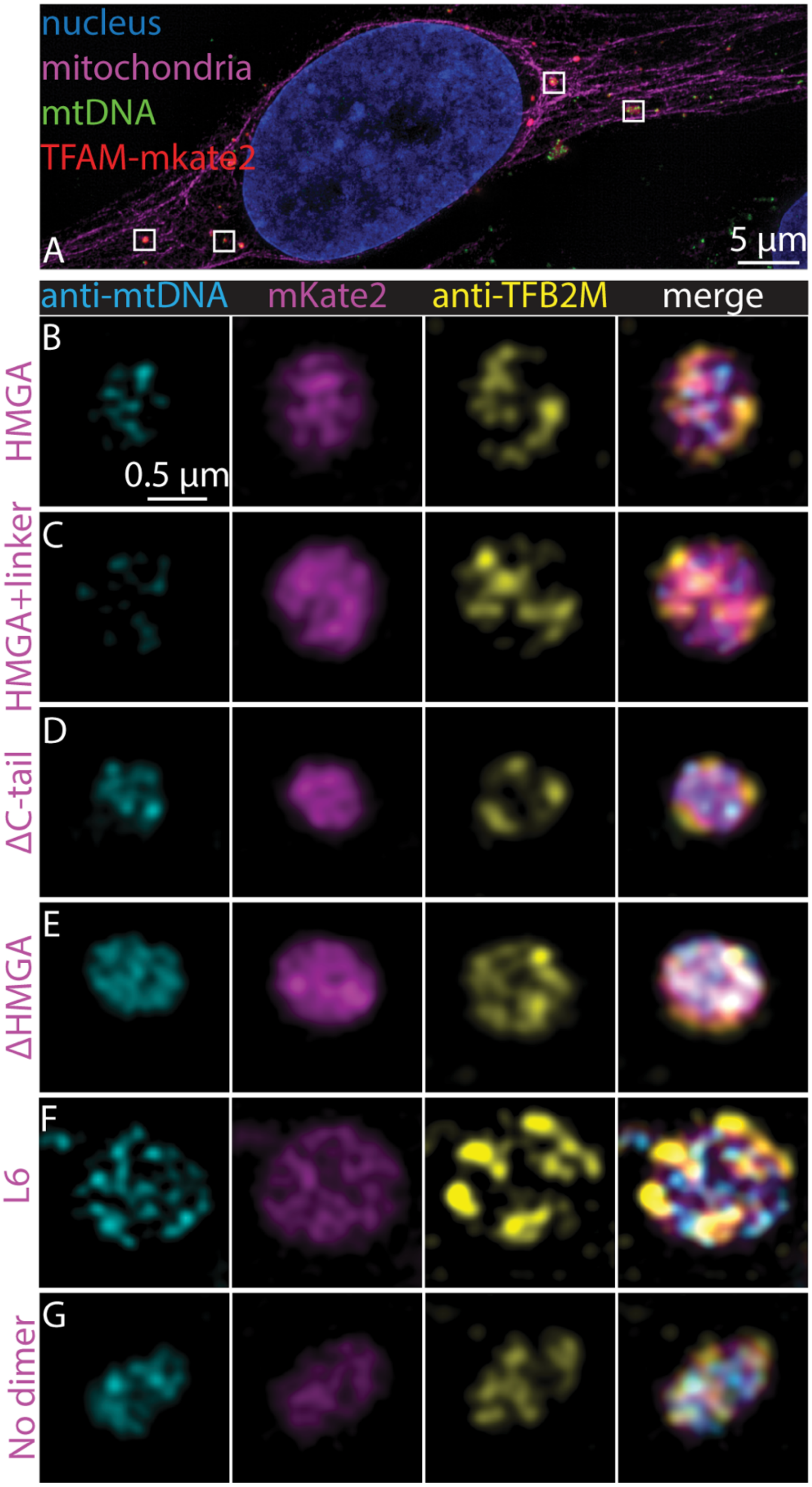

**Fig. S5**

(A) SIM maximum intensity projection of a fixed HeLa with the nucleus (DAPI, blue), mitochondria (MitoTracker Deep Red, magenta), mtDNA (anti-DNA, green), and TFAM (TFAM-mKate2, red) from which nucleoids were shown in Fig. 5A-C. Scale bar = 5 µm. White boxes indicate nucleoids that were chosen for panels in A-C and in Fig. 5. (B-G) Single z-slices of SIM images of nucleoids after over-expression of mutants in HeLa cells, including HMGA-mKate2 (B), HMGA+linker-mKate2 (C), ΔC-tail-mKate2 (D), ΔHMGA-mKate2 (E), L6-mKate2 (F), and no-dimer-mKate2 (G). Left most panel is of mtDNA (anti-DNA, cyan), followed by the mutant (mKate2, magenta), TFB2M (anti-TFB2M, yellow), and merged images. Scale bar = 0.5 µm.

**Supplementary Figure S6:**

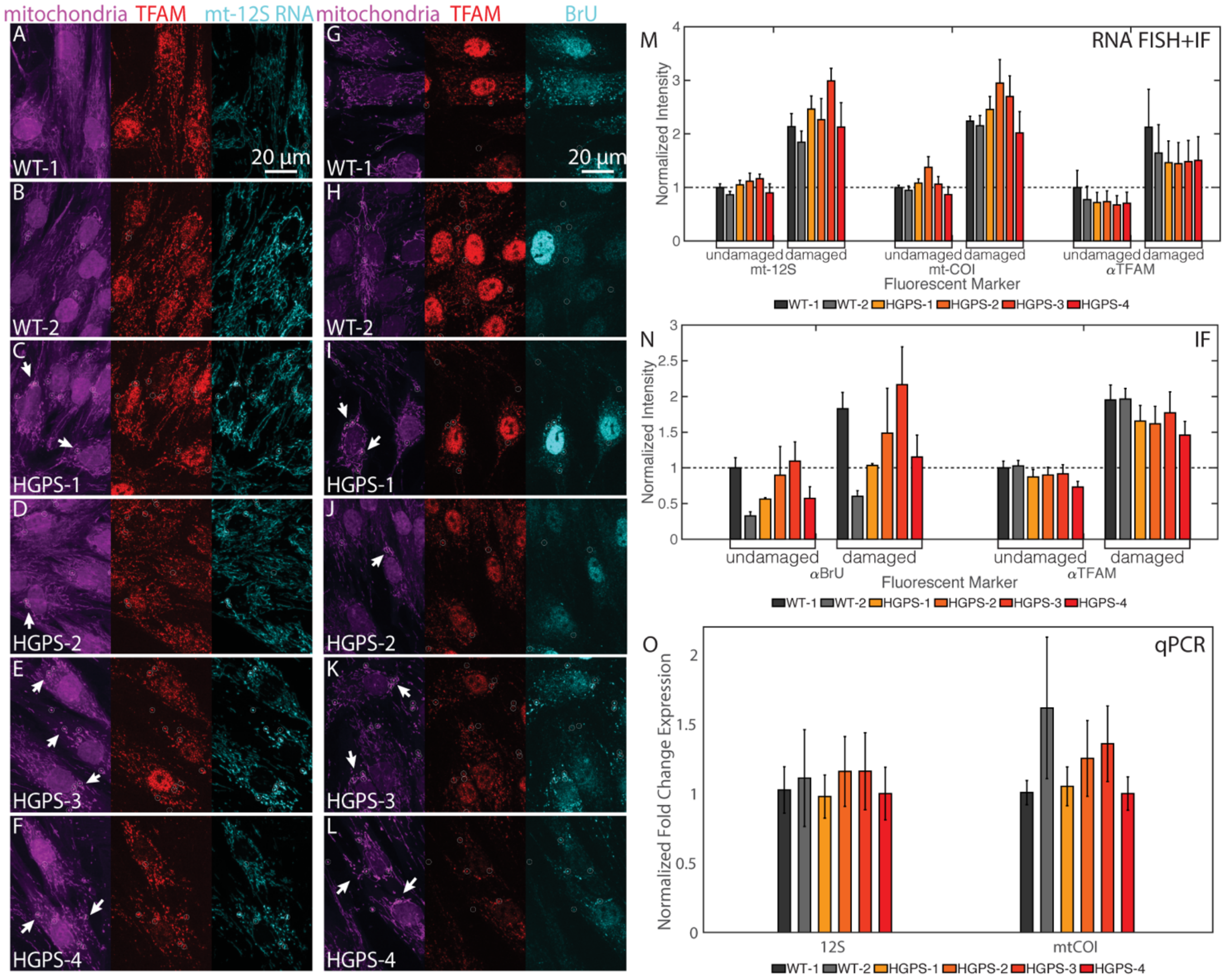

**Fig. S6:** (A-F) Representative high-throughput confocal images of WT-1 (A), WT-2 (B), HGPS-1 (C), HGPS-2 (D), HGPS-3 (E), and HGPS-4 (F) cell lines. Left image is mitochondria (MitoTracker Red, magenta); center image is TFAM (anti-TFAM, red); right image is mt-12S rRNA (RNA FISH, red). Scale bar = 20 μm. (G-L) Representative high-throughput confocal images of WT-1 (G), WT-2 (H), HGPS-1 (I), HGPS-2 (J), HGPS-3 (K), and HGPS-4 (L) cell lines. Left image is mitochondria (MitoTracker Red, magenta); center image is TFAM (anti-TFAM, red); right image is of nascent transcripts after BrU incorporation (anti-BrdU, red). Scale bar = 20 μm. (M-N) Based on high-throughput imaging, integrated intensity values for markers for mt-12s rRNA and mt-COI RNA (M) or BrU (N) along with anti-TFAM (M-N) were compared for nucleoids from undamaged and damaged (indicated by white circles in earlier images) mitochondria per cell and averaged, where n=3 experimental replicates and error bars represent standard error. (O) qPCR results for mitochondrial transcripts 12S and mt-COI for all cell lines. n=3 independent experimental replicates, each with three technical replicates, and error bars represent standard error.

**Supplementary Videos**

**Video S1: Fusion of nucleoids under phototoxic conditions in HGPS cells.** Live primary HGPS cells were incubated with MitoTracker Deep Red (mitochondria, magenta) and PicoGreen (mtDNA, green) prior to imaging with a laser scanning confocal (LSM780) microscope. Maximum Intensity Projection (MIP) of z-stacks acquired every 30 seconds is shown. Scale bar = 2 microns.

**Video S2: Fusion of nucleoids under phototoxic conditions in WT cells.** Live primary WT cells were incubated with MitoTracker Deep Red (mitochondria, magenta) and PicoGreen (mtDNA, green) prior to imaging with a laser scanning confocal (LSM780) microscope. Maximum Intensity Projection (MIP) of z-stacks acquired every 30 seconds is shown. Scale bar = 2 microns.

**Video S3: Fusion of nucleoids under phototoxic conditions and with EtBr in HGPS cells.** Live primary HGPS cells were incubated with MitoTracker Deep Red (mitochondria, magenta) and PicoGreen (mtDNA, green) along with Ethidium Bromide for ~30 minutes prior to imaging with a laser scanning confocal (LSM780) microscope. Maximum Intensity Projection (MIP) of z-stacks acquired every 30 seconds is shown. Scale bar = 2 microns.

**Video S4: Fusion of nucleoids under phototoxic conditions and with EtBr in WT cells.** Live primary WT cells were incubated with MitoTracker Deep Red (mitochondria, magenta) and PicoGreen (mtDNA, green) along with Ethidium Bromide for ~30 minutes prior to imaging with a laser scanning confocal (LSM780) microscope. Maximum Intensity Projection (MIP) of z-stacks acquired every 30 seconds is shown. Scale bar = 2 microns.

**Video S5: FRAP of TFAM droplet *in vitro*.** FRAP on droplet produced at 25 μM TFAM, 150 mM NaCl, 20 mM Tris-HCl, pH 7.5, 30 minutes after mixing. Time interval = 15 s. Scale bar = 1 micron.

**Video S6: FRAP of TFAM-mtDNA droplet *in vitro*.** FRAP on droplet produced at 25 μM TFAM, 100 ng/µl mtDNA, 150 mM NaCl, 20 mM Tris-HCl, pH 7.5, 30 minutes after mixing. Time interval = 15 s. Scale bar = 1 micron.

**Video S7: FRAP of single small nucleoid in live cells.** An entire nucleoid was bleached in live HeLa cells expressing TFAM-mKate2 (red) and labelled with PicoGreen (green). Time interval = 10 s. Scale bar = 1 micron.

**Video S8: FRAP of single small nucleoid followed by fusion in live cells.** An entire nucleoid was bleached in live HeLa cells expressing TFAM-mKate2 (red) and labelled with PicoGreen (green) followed by a liquidlike fusion event with a neighboring nucleoid. Time interval = 10 s. Scale bar = 1 micron.

**Video S9: FRAP of a portion of large nucleoid in live cells.** Partial bleach of an enlarged nucleoid in live HeLa cells expressing TFAM-mKate2 (red) and labelled with PicoGreen (green). Time interval = 25 s. Scale bar = 1 micron.

**Video S10: Fusion of nucleoids under phototoxic conditions and with EtBr in HeLa cells.** Live HeLa cells were incubated with Ethidium Bromide for ~30 minutes prior to imaging with laser scanning confocal (LSM780) microscope. HeLa cells over-expressing TFAM-mKate2 (red) were labelled with PicoGreen (green) to visualize mtDNA and with MitoTracker Deep Red (magenta) to visualize mitochondria. MIP of z-stacks acquired every two minutes is shown. Scale bar = 2 microns.
